# Differential contribution of the RNA recognition motifs to SRSF1 protein interaction network

**DOI:** 10.1101/2025.11.17.688470

**Authors:** Pablo Mammi, Andrea Pawellek, Nicolás Gaioli, Laureano Bragado, Berta Pozzi, Hernan Grecco, Angus I. Lamond, Anabella Srebrow

## Abstract

The protein SRSF1, initially identified as a splicing factor, is currently known to participate in a variety of cellular processes. However, its role and regulatory mechanisms in these processes remain incompletely understood. Using a SILAC proteomic approach, we have characterised the SRSF1 interactome, thereby identifying multiple diverse protein interaction partners of SRSF1 and the molecular complexes with which it is associated. Given that the RNA-binding ability of SRSF1 is necessary for many of its activities, we extended the analysis of the SRSF1 interactome to compare the wild type protein against two site-specific SRSF1 variants, bearing mutations in each of its two RNA recognition motifs (RRMs). The first mutation disrupted the canonical RRM (RRM1), while the second affected the pseudo-RRM (RRM2). Interestingly, the data show clear differences between the SRSF1 interacting proteins, depending on the functional status of each RRM domain. In particular, a functional RRM1 was critical for maintaining interactions with multiple partners, especially those involved in translation-related processes, showing a far greater loss of interactions upon mutation than RRM2. This study provides valuable new insights into the broader cellular roles and interaction landscape of SRSF1.

## INTRODUCTION

The Serine/arginine-rich splicing factor 1 (SRSF1), previously named SF2/ASF, is largely known as a key regulator of pre-mRNA processing [1], [2]. While SRSF1 is not a component of the core snRNP spliceosome subunits, its dual role as both a constitutive and alternative splicing factor has been widely described and SRSF1 is now seen as an archetypal auxiliary component of the splicing machinery. Nevertheless, splicing is likely not the only cellular process in which SRSF1 is involved; over the years, a diversity of other potential functions has been associated with this protein [3].

SRSF1 is a prototypical member of the SR protein family. All SR proteins share a conserved modular structure comprising either one, or two N-terminal RNA-recognition motifs (RRMs) and a C-terminal domain of variable length that is rich in arginine and serine dipeptides (RS domain) [4]. Different SR protein family members are involved in a range of cellular processes, including gene expression, RNA processing and translation [5], [6]. Although these proteins are mainly found in the nucleus, many can shuttle between the nucleus and the cytoplasm, at different rates [7], [8]. With respect to functions and cellular pathways beyond splicing [3], [9], SRSF1 has been reported to be involved in: additional aspects of mRNA metabolism, such as stability/degradation [10] and mRNA export [11]; stimulation of transcriptional elongation [12]; genome stability [13]; translation regulation [14], [15]; nonsense-mediated decay (NMD) [16], [17]; regulation of protein post-translational modification by SUMO conjugation [18], [19]; and pri-miRNA processing [20], [21], [22]. Although the regulatory functions of SRSF1 in many of these processes appear to be independent from its role as a splicing factor, mutations affecting its RNA binding ability disrupt its activity in several of these pathways [23].

Even though RS domains have been shown to directly interact with RNA molecules [24], [25], most SR proteins, including SRSF1, bind RNA via one or more RRMs, like the general RNA-binding proteins (RBPs). SRSF1 has an N-terminal RNA-binding domain, called RRM1. This, and other canonical RNA binding domains, typically assume a β_1_α_1_β_2_β_3_α_2_β_4_ topology. They are characterized by having two sub-motifs, termed RNP-1 and RNP-2 (located within β_3_ and β_1_ respectively, in antiparallel orientation), with aromatic and positively charged amino acids, which constitute the RNA binding site [26], [27], [28], [29]. The second RNA recognition motif in SRSF1 is RRM2, a non-canonical, or pseudo-RRM. Unlike the canonical ones, pseudo-RRMs do not contain aromatic amino acids responsible for binding to RNA within their β-sheets, suggesting an unconventional mode of interaction. Some studies have pointed to a conserved heptapeptide (SWQDLKD), centred on an α_1_-helix, as responsible for the RNA binding ability[26], [30], [31].

For the characterization of the multiple functions of SRSF1, the disruption of the RRMs has been a powerful approach that allows studying the role it plays in particular cellular processes as an RNA-binding protein, as well as the molecular basis underlying each of its tasks. Particularly, mutational analyses have shown specific residues within either SRSF1 RRM1, or RRM2, that are necessary for RNA binding. This includes phenylalanines 56 and 58, within the RNP-1 submotif of RRM1 [32] and tryptophan 134 of the conserved heptapeptide within RRM2 [11], [31], [33]. The substitution of these amino acids not only interferes with the RNA binding ability of each RRM [11], [32], but also affects the splicing activity of SRSF1 [31], [32], [34]. In this context, two SRSF1 variants have been widely used, due to their ability to abolish the RNA binding activity of these domains. The first contains a double substitution of phenylalanines 56 and 58 with aspartic acid and is termed SRSF1-(FFDD). The second contains a single point mutation where tryptophan 134 is replaced by arginine and is known as SRSF1-(W134A) (Figure 1A). The inability of both mutants to bind RNA has been confirmed *in vitro* by UV-crosslinking experiments [11], [32]. Over time, several activities of SRSF1 have been tested with these two variants as RNA-binding mutants, but they have been rarely used together in the same study [34], [35]. The most relevant findings achieved with these mutants are summarized below.

**Figure 1:**
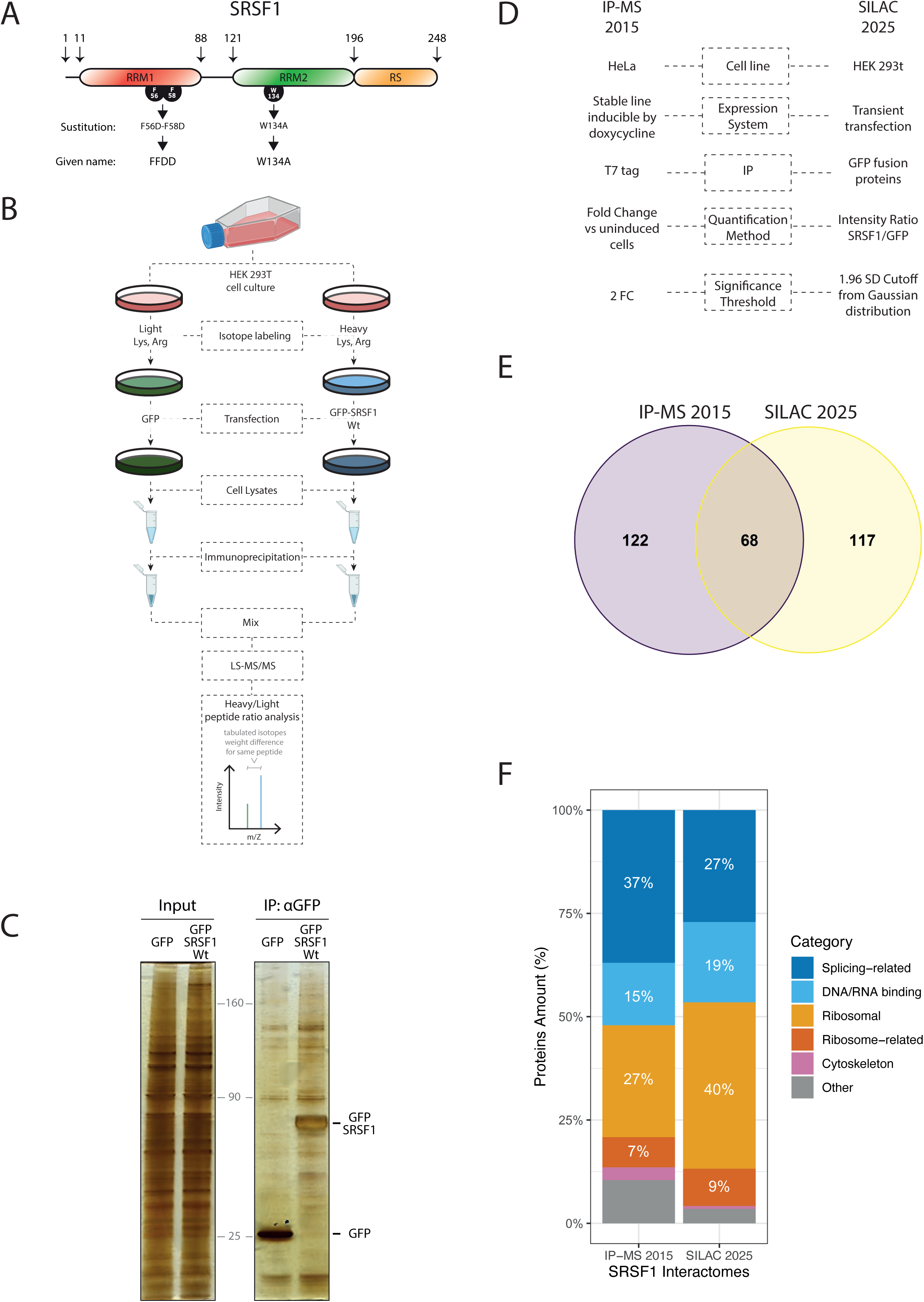
SRSF1 interactome analysis. (A) Schematic representation of the modular structure of SRSF1, depicting the three functional domains. Highlighted residues display the one-letter code and position of key amino acids within RNA recognition motifs (RRMs), with annotations specifying the substitutions introduced in the mutant variants used as well as the name by which we refer to them in this work. Black arrows indicate the position of selected amino acids. (B) Schematic representation of the experimental design for the SILAC assay. A unique cell culture was split into two, each incubated with distinct isotopically labeled amino acids over several passages to ensure complete incorporation. Cells were then transfected with expression plasmids for GFP or GFP-SRSF1 Wt and lysed 24 hours after transfection. Cell lysates were subjected to GFP immunoprecipitation, and the resulting immunoprecipitates were combined. Finally, samples were analyzed by mass spectrometry to quantify the ratio of peptides labeled with one isotope (heavy) relative to those labeled with the other (light). (C) Comparison between a representative sample of immunoprecipitates from GFP alone- and GFP-SRSF1-expressing cells, monitored by SDS-page and subsequent silver staining. (D) Comparison of the experimental conditions of the interactome carried out by Akerman *et al*. in 2015 and our current SILAC. (E) Number of proteins enriched in SRSF1-immunoprecipitated samples. The yellow area indicates significant hits in the SILAC experiment. The purple area indicates significant hits in the IP-MS performed by Akerman *et al*. The merge indicates common hits detected by both approaches. (F) Results of the interactomes analysis. The major functional categories of proteins identified as SRSF1 interactors are depicted. The white number indicates the percentage value of the corresponding category with respect to the total set of interactors.

FFDD: This is the most widely used RNA-binding-deficient mutant of SRSF1. The founding publication showed using *in vitro* assays that it is unable to function in constitutive splicing [32]. Multiple subsequent studies demonstrated that SRSF1-(FFDD) fails to carry out several of the activities associated to the wild type protein: SRSF1-(FFDD) cannot stimulate translation [36]; it has lost the ability to complement cell survival in SRSF1-depleted cells [37]; fails to bind to exonic splicing enhancers (ESE) [34], [38]; does not mediate mRNA instability [39]; fails to promote tumorigenesis by enhancing β-catenin biosynthesis [40]; and is unable to regulate pri-miRNA biogenesis [20]. In addition, it shows a reduced interaction with the long non-coding RNA MALAT1 [41]. While this mutant is unable to shuttle between the nucleus and the cytoplasm [8], [35], displaying predominantly nuclear localization, it still accumulates in splicing factor reservoirs, termed speckles, and it is recruited to transcriptionally active sites [42].

W134A: This variant was more recently described [11]. In addition to its inability to bind RNA, it was also shown to lack the ability to stimulate translation [11]. In 2010, it was reported that, like the wild type protein, SRSF1-(W134A) predominantly localizes to the nucleus at steady state but is unable to shuttle between the nucleus and the cytoplasm [35]. Similar to the SRSF1-(FFDD) mutant, SRSF1-(W134A) is highly defective in ESE binding [34]. Some studies have also employed an SRSF1 variant, referred to as AAA, in which the WQD residues in the conserved heptapeptide are replaced by three alanines. This mutant, which includes the W134A substitution, is non-functional in promoting β-catenin accumulation in tumorigenesis [40]; does not bind to either PP2A or mTOR and therefore fails to regulate eIF4E-dependent translation [15]; and it is not recruited to nuclear stress bodies [43].

The results obtained with these mutants suggest that SRSF1 depends on its RNA binding ability for multiple functional roles. However, the impact of the disruption of RNA binding by each RRM in those processes has not been fully characterised. Since the multifaceted properties of SRSF1 are a consequence not only of its RNA-binding ability, but also of its nuclear-cytoplasmic shuttling as well as its involvement in different multiprotein complexes [44], the interpretation of the molecular mechanisms underlying SRSF1 multiple roles can be intricate and challenging.

To advance our understanding of the multifunctional SRSF1 protein and the role of RNA binding in its various activities, we set out to identify the SRSF1 interactome and to interrogate the contribution of each RRM to the SRSF1 interaction network. To this end, we employed mass spectrometry (MS)-based proteomic analysis, using SILAC [45], to compare the co-immunoprecipitated protein interaction partners associated, respectively, with GFP-tagged wild type SRSF1 and with each of the RNA binding-defective mutants, SRSF1-(FFDD) and SRSF1-(W134A). The data show that SRSF1 wild type interacts not only with splicing factors but also with multiple other cellular proteins. Furthermore, the data show that many of the observed interactions are lost upon mutations in SRSF1 that abrogate RNA binding, via either RRM1, or RRM2.

## RESULTS

### SRSF1 Interactome

The splicing factor SRSF1 is involved in a variety of cellular processes, carried out through direct or indirect interactions within intricate protein or ribonucleoprotein complexes. To study the protein interaction partners of SRSF1, we used a quantitative, mass spectrometry-dependent MAP-SILAC proteomic approach to analyse cells expressing GFP-tagged SRSF1 wild type [46], [47]. For this purpose, HEK 293T cells were grown at 37 °C in media containing either light, or heavy isotopic forms of lysine and arginine. Light-labelled cells were transfected with a GFP expression vector as a control, while heavy-labelled cells were transfected with a GFP-SRSF1 wild type (Wt) expression vector. At 24 h post transfection, an equal number of cells from each SILAC channel were used to generate whole cell lysates. These were subjected to immunoprecipitation with an anti-GFP nanobody coupled to magnetic beads (GFP-trap®), and the precipitated samples were combined and prepared for LS-MS/MS analysis (Figure 1B). GFP immunoprecipitation and the presence of specific SRSF1 binding partners was observed by comparing the immunoprecipitates on a silver-stained SDS-page (Figure 1C). This experiment was repeated with three biological replicates. For the evaluation of mass spectrometry results, from the total peptide intensity data, those protein-groups recognized by at least two unique peptides were considered for further analysis. High-confidence enriched proteins in GFP-SRSF1 immunoprecipitates were identified by those Heavy/Light (H/L) intensity ratios that exceeded the threshold of significance established in each replicate (see Materials and Methods).

Based on these considerations, we identified 185 proteins that specifically co-immunoprecipitated with GFP-SRSF1 Wt (Supplemental Table 1). Of these proteins, 23 correspond to known interactors of SRSF1, according to the interaction repository BioGRID [48] (highlighted rows in Supplemental Table 1). Among those, several are well-known direct partners, such as other SR proteins (SRSF3, SRSF7, SRSF10); SR protein-specific kinases (SRPK1, SRPK2, SRPK3); heterogeneous nuclear ribonucleoproteins (hnRNP A0, hnRNP A1, hnRNP F, hnRNP U); and the spliceosome protein U1A 70k. Notably, some previously reported interactors were not detected above our SILAC significance threshold in this dataset, which may reflect differences in protein abundance or interaction dynamics under the specific cellular conditions analysed here. Otherwise, a variety of hitherto non-reported interactors were detected in our study, some of which could represent novel SRSF1 direct partners awaiting validation, while others may indirectly interact with SRSF1 within large macromolecular complexes. The latter could be the case for multiple ribosomal proteins, taking into account the already described regulatory role of SRSF1 in protein translation and its co-sedimentation with ribosomal particles [15], [36].

### Comparative proteomics of SRSF1 reveals its involvement in diverse molecular complexes

In 2015, Akerman *et al*. investigated the protein connectivity of splicing factors using probabilistic network reconstruction [49]. Employing SRSF1 as a prototypical SR protein, their proposed model was validated by immunoprecipitation and mass spectrometry. We decided to compare our SILAC results with this already available interactome data, in order to test the robustness of our proteomic experimental design to elucidate the SRSF1 association landscape.

Despite the differences between the two experimental settings (Figure 1D), of the total list of significantly co-immunoprecipitated proteins, 68 were common hits to both proteomics approaches (Supplemental Table 1), representing the 37% of our validated proteins (Figure 1E).

In addition, Akerman *et al.* demonstrated that nearly half (56%) of the total interactions they had detected were resistant to nuclease treatment. Considering the 68 hits that are common to both studies, only 17 interactions (25%) can be classified as nucleic acid independent, according to Akerman *et al.* (Supplemental Table 1). This lower percentage, with respect to Akerman’s approach, allow us to estimate by extrapolation that 3 out of 4 interactions detected in our study could be mediated by RNA, which highlight the strength of using SILAC to detect not only direct or indirect protein-protein interactions, but also those occurring within ribonucleoprotein complexes.

To get a better understanding of the composition of these multiple molecular complexes, we proceeded to classify the proteins enriched in SRSF1-immunoprecipitates into six categories: (i) Splicing-related, (ii) DNA/RNA binding, (iii) Ribosomal, (iv) Ribosome-related, (v) Cytoskeleton and(vi) Others. As expected, almost a half of the detected hits are Splicing-related or DNA/RNA binding proteins (Figure 1F), reflecting the already known nature of SRSF1 as an essential splicing factor and an RNA binding protein. Consistent with this, KEGG pathway enrichment analysis of the same dataset identified the spliceosome and mRNA surveillance as the most significantly enriched pathways (Supplemental Figure 1A). Remarkably, a similar percentage of identified proteins are Translation-related (Ribosomal and Ribosome-related). Knowing that most, although not all, of those latter interactions are nuclease-treatment sensitive, according to Akerman and collaborators [49], and that ribosomal proteins are among the most abundant within the cell, we hypothesise that these categories may represent proteins that immunoprecipitated in association with translationally active SRSF1-bound mRNAs. Nevertheless, taking into account the known involvement of SRSF1 in translation regulation, we cannot rule out the possibility that many of those interactions are also direct. Remarkably, when we compared our proteomic dataset with that of Akerman *et al*., the ribosome emerged as the most enriched KEGG pathway among the shared hits (Supplemental Figure 1B), suggesting that SRSF1 translation-related interactions are robust and reproducible across distinct experimental approaches.

The category analysis shown in Figure 1F yielded similar outcomes for the SILAC and IP-MS methods, demonstrating the consistency of both experimental strategies. Also, it reflects the diverse molecular complexes in which SRSF1 is involved, many of which are due to its RNA-binding ability, as shown by Akerman *et al*., and to the fact that this protein accompanies mRNA molecules along different steps of their nucleus-cytoplasmic metabolism and transport journey, being part of messenger ribonucleoprotein particles (mRNPs) [50]. This varied repertoire of macromolecular interactions is thus in agreement with the multitasking nature of SRSF1 [23], [51], [52].

### Characterization of SRSF1 RNA-recognition motif mutants with respect to their sub-cellular localization and splicing

To understand the contribution of the RNA-recognition motifs (RRMs) to the association landscape of SRSF1, we chose to work with two already described and widely used variants, obtained by point mutations disrupting either RRM1- (FFDD mutant) or RRM2- (W134A mutant) mediated SRSF1 RNA-binding ability. Both mutants were designed so as not to disrupt the proper folding of the protein [11], [32], and AlphaFold predicts identical structures for the mutant RRM domains compared to the Wt domain (Supplemental Figure 2). We therefore focused on comparing their functional behavior, to assess whether major differences emerge despite the absence of detectable structural alterations.

As a first approach, we explored the cellular distribution of each variant to ensure that this could not be the cause of large differences in their involvement in cellular mechanisms. For all SRSF1 versions evaluated in this study, i.e., Wt, FFDD and W134A mutants, a predominantly nuclear localization was reported [8], [35]. To analyse possible changes in the subcellular localization of the RNA binding mutants, we performed two microscopy-based experiments. Splicing factors, and in particular SRSF1, are stored in nuclear interchromatin granule clusters, usually termed speckles [53]. Thus, we first assessed a putative differential accumulation of any SRSF1 variants in speckles. Upon overexpressing GFP-fused versions of SRSF1 Wt, W134A and FFDD in HeLa cells, fluorescence intensity in nuclear speckles was quantified using confocal microscopy images. Both mutants exhibit significantly (p < 0.05) reduced nuclear speckle localization at steady state, compared with Wt (Figure 2A).

**Figure 2:**
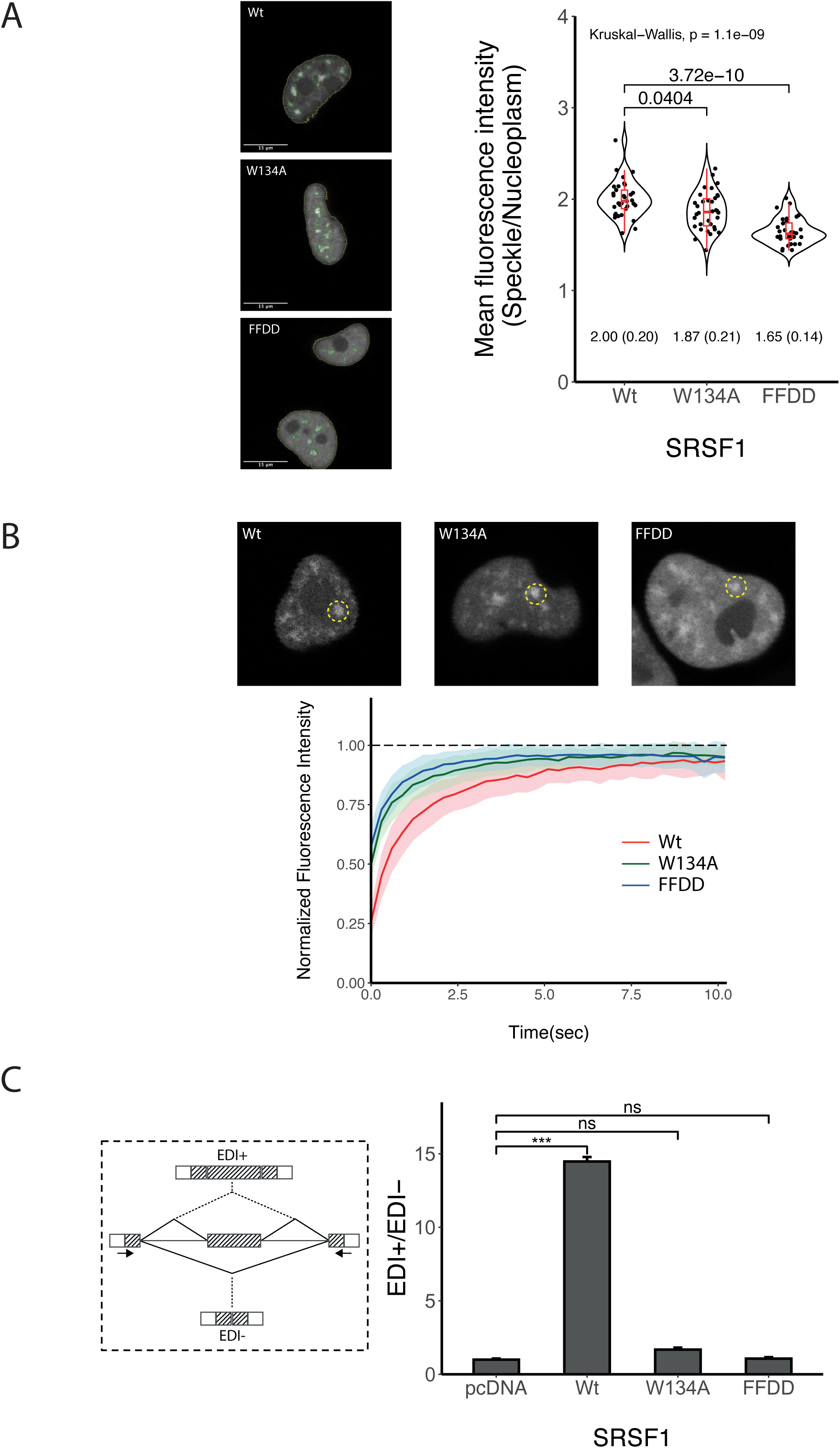
Subcellular localization and alternative splicing activity of SRSF1 RNA-binding mutants. (A) Nuclear speckle localization of SRSF1 variants. HeLa cells were transfected with the indicated GFP-SRSF1 variants. After 24 h, cells were fixed and mounted. Images taken by confocal microscopy were analyzed with *CellProfiler* software. A pipeline was developed to automatically identify nuclei and speckles in each cell. A representative photo of each mutant is shown in the left panel. The yellow lines delimit the nuclei while the green lines the speckles. The mean intensities of the speckles and nucleoplasm in 30 cells were quantified, and the ratio between these two measurements was plotted as a dot for each cell. The violin graph reflects dots density, where the central red line within the box-plot corresponds to the median of the plotted data. The brackets indicate the between-group p-value of the Dunn’s post-hoc test for multiple comparisons. The mean of the plotted values and the corresponding standard errors (within parenthesis) are indicated below each violin-plot. (B) Dynamics of fluorescence recovery in speckles. FRAP analysis shows differences in post-photobleaching recovery of both SRSF1 mutants with respect to the Wt version. HeLa cells transiently transfected with GFP-SRSF1 variants were used for this experiment. A representative cell from each group is displayed in the upper panel. The yellow circle indicates the area of the speckle where the laser pulse was applied and where the fluorescence was measured before and after this event. Fluorescence intensity is expressed relative to pre-photobleaching fluorescence. Each curve is the average of the intensity measured for 25 to 30 cells. The shaded areas in color that accompany each curve represent the standard deviation for the values of each group. (C) HeLa cells were co-transfected with the splicing reporter minigene and expression vectors for different SRSF1 variants or pcDNA empty vector. After 24 h, total RNA was prepared from whole cell lysates, and used for reverse transcription (RT) followed by radioactive end-point PCR. PCR products were run on native polyacrylamide gels, which were dried and exposed to radiographic plates (representative autoradiographs are shown in Supplemental Figure 3C). The bands corresponding to each isoform (EDI+ or EDI-) were cut and the radioactivity was measured in a scintillation counter. The quantification of three biological replicates is shown in a bar graph, where the ratio between the inclusion mRNA isoform over the exclusion mRNA isoform is observed. The brackets indicate the between-group p-value determined using a paired two-tailed t-test. Significant p-values are indicated by the asterisks above the graphs (***P < 0.001; **P < 0.01; *P < 0.05). Scheme of the alternative splicing reporter mini-gene is on the left box. The rectangles represent exons and the lines introns. The striped exons correspond to the fibronectin gene, whereas the white exons to the α-globin gene. In the center of the diagram the region of interest contained within the plasmid is observed while in the upper part the mRNA isoform containing the alternative exon EDI (mRNA EDI +) derived from the mini-gene is shown, and in the lower part the mRNA isoform lacking the alternative exon or exclusion isoform (mRNA EDI -). PCR primers used are indicated by the black arrows spanning the junctions of the hybrid exons (fibronectin-α-globin).

Second, to get further insight into the dynamics of SRSF1 association/dissociation with speckles, a Fluorescence Recovery After Photobleaching (FRAP) experiment was performed. The two RRM mutants showed a higher rate of fluorescence recovery in speckles than SRSF1 Wt (Figure 2B). This mobility depends on both diffusion and potential binding interactions. A deeper analysis revealed that not only the diffusion coefficient, but also the mobile fraction of both mutants shows lower values compared to the Wt version of this protein (Supplemental Figure 3A). These results could be indicative of an extremely fast initial recovery of intensity by the FFDD and W134A mutants, in comparison with Wt SRSF1, denoting that RNA-binding mutants were retained at speckles for a shorter period of time, in agreement to what has been shown in Figure 2B. Microscopy results hint that, despite the fact both RRM mutants accumulate in speckles inside the nucleus, they lose interaction with components of these sub-nuclear compartments, decreasing their residence time in these granules.

Having ruled out extreme changes in subcellular localization, we next assessed their role as alternative splicing regulators, the prototypical activity of SRSF1. To this end, the effect of overexpressing each SRSF1 variant on fibronectin alternative splicing was monitored, by analysing mRNA isoforms (either containing or lacking the fibronectin alternative exon EDI) derived from a widely used splicing reporter minigene (scheme on Figure 2C) [54], [55]. Cells were co-transfected with the FN EDI splicing reporter and either the expression vector for T7-SRSF1 Wt, T7-SRSF1 (W134A), T7-SRSF1 (FFDD), or an empty pcDNA plasmid control (Supplemental Figure 3B). While overexpression of Wt SRSF1 stimulates EDI exon inclusion, as previously reported [56], neither the W134A, nor FFDD mutant, exerted any effect on this alternative splicing event in these experimental conditions (Figure 2C, Supplemental Figure 3C).

### SRSF1 variants reveal differential contribution of RRM domains to the protein interaction network

To decipher the contribution of each RRM to SRSF1 interactome, we performed a second SILAC-MS co-IP experiment. To this end, HEK 293T cultured cells were labelled with light (L), medium (M) or heavy (H) stable-isotopes of lysine and arginine, corresponding to cells transiently expressing either SRSF1 Wt, or, respectively, the SRSF1 W134A or FFDD mutants. Whole cell lysates from each condition were prepared and processed for analysis by co-immunoprecipitation MS, as described above (Supplemental Figure 4A).

From the proteins identified that co-precipitated with GFP-SRSF1, we focused on those that are either absent, or reduced in abundance, in the immunoprecipitates of the SRSF1 mutants, compared with Wt. The resulting proteomic data show the mean isotope ratio of SRSF1 Wt vs mutant for all quantifiable peptides, plotted for every protein group (Figure 3A). For ease of comparison, the ratio is shown as a positive value for SRSF1-(W134A) and as a negative value for SRSF1-(FFDD). From these data it is evident that, in comparison with SRSF1 Wt, SRSF1-(FFDD) co-immunoprecipitates with fewer interactors than SRSF1-(W134A), indicating that the SRSF1-(FFDD) mutant interacts stably with less partner proteins than SRSF1-(W134A). This is already detectable in a silver stained SDS-page gel from samples co-precipitated with the respective GFP-SRSF1 variants (Supplemental Figure 4B).

**Figure 3:**
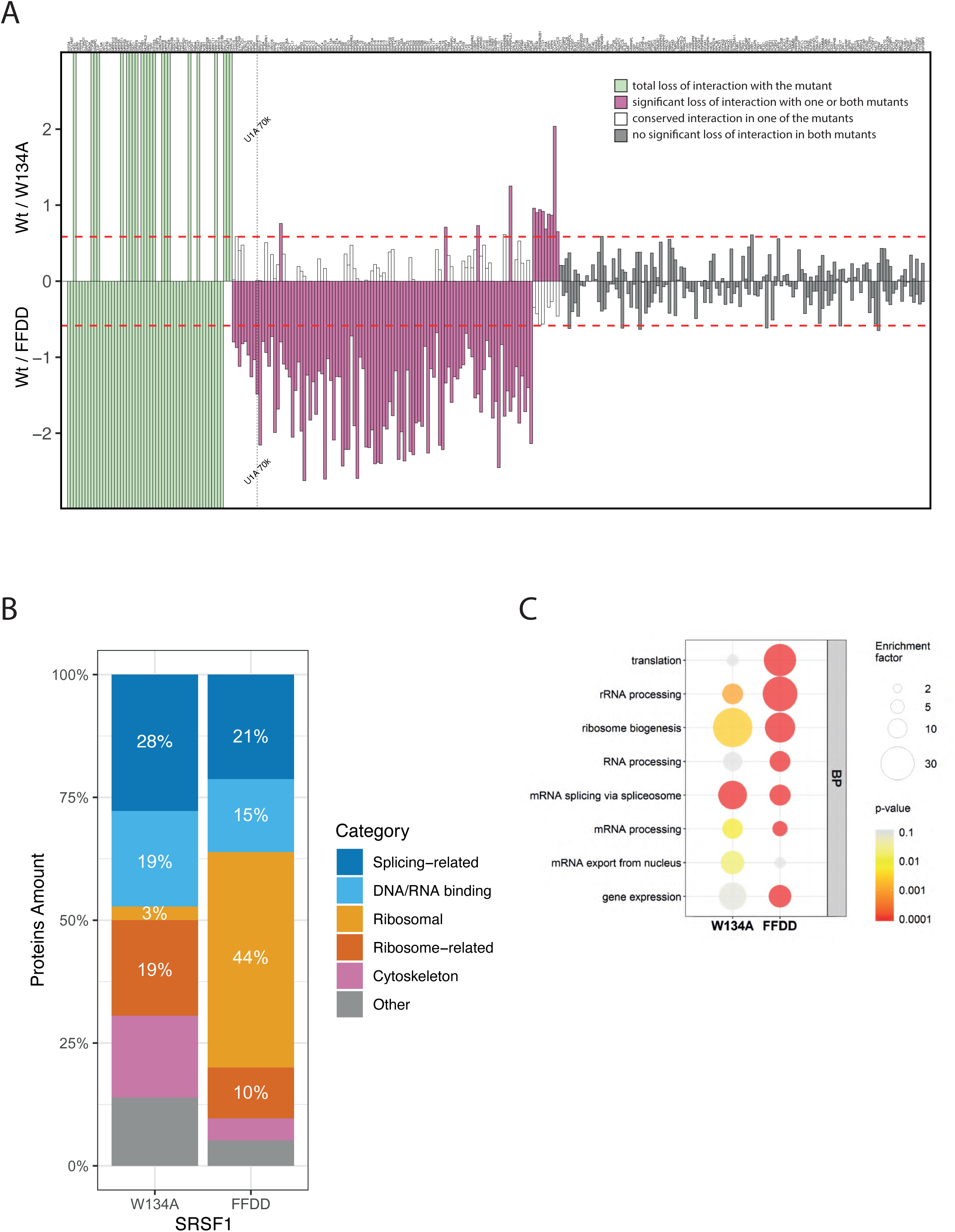
SILAC analysis of SRSF1 RNA-binding mutants. Three HEK 293T cultures derived from a common batch were each incubated with a different stable isotope-labeled version of two amino acids. Each culture was then transfected with expression vectors for GFP-SRSF1 Wt, GFP-SRSF1 W134A or GFP-SRSF1 FFDD. After 24 h, whole cell lysates were subjected to immunoprecipitation with GFP-trap and subsequently prepared for mass spectrometry analysis. The peptide ratios between each of the conditions were calculated to quantify the relative abundance of these in each immunoprecipitate. We focus on those proteins with which the mutants lose interaction compared to SRSF1 Wt. (A) Bar plot of the ratios between the intensity of quantifiable peptides in the immunoprecipitates of SRSF1 Wt over those of each RRM mutant. The values are expressed on a logarithmic scale with base 2, and for easy comparison as positive values for the ratios Wt/W134A and negative values for those of Wt/FFDD. Green bars indicate proteins that appear only in SRSF1 Wt immunoprocipitates and for which no quantifiable peptides are found with the mutant. The purple bars indicate proteins with which this mutant significantly loses interaction with respect to Wt. The white bars indicate that there are no significant differences for that particular protein in the corresponding mutant, but there are for the other RRM-mutant. The grey bars indicate that no significant differences were found for either of the two mutants. The red dashed lines mark 1.5-fold-enrichment in immunoprecipitates of Wt with respect to the mutants (although these differences may not be significant due to experimental variability). The name of the protein according to the Uniprot code is indicated above each bar. Data obtained for the U1-70K protein are highlighted with a vertical dashed line. The Excel file with the data of this plot is available upon request. (B) Results of the interactome analysis. The main functional categories of proteins identified as losers of interaction with the RRM mutant relative to the SRSF1 Wt are described. The white number indicates the percentage value of the corresponding category. (C) GO analysis of proteins showing reduced interaction with SRSF1 mutants compared to the wild-type protein. The diameter of the circles represents the enrichment factor of these groups, and the color indicates the p-value of the statistical analysis, according to the legend on the right panel. (BP) Biological processes.

Beyond differences in the number of proteins that lose interaction with each mutant as compared with SRSF1 Wt (green and purple columns in Figure 3A), the identity of these proteins is also of particular interest. Based on GO term analysis, no major distinctions were detected between the molecular functions of the diminished interactors identified in the immunoprecipitates of the FFDD mutant and those of the W134A mutant (Supplemental Figure 4C). To better compare the interaction profiles of the SRSF1 mutants we categorized the proteins showing reduced interaction with each mutant, relative to the Wt, into the six functional groups previously defined, and plotted their relative proportions (Figure 3B).

In the case of mutant W134A, the proportions of splicing-related proteins and RNA/DNA-binding proteins among the set of interacting proteins seen with Wt but not with this mutant (28% and 19%, respectively) closely match their representation within the total Wt SRSF1 interactome shown in Figure 1D (27% and 19%). This suggests that the W134A mutation does not selectively affect SRSF1 interaction with RNA/DNA-binding proteins, but rather that the losses reflect the overall composition of the Wt interactome. In contrast, the FFDD mutant loses a lower proportion of interactors from these two categories (21% and 15%, respectively). Interestingly, a major difference emerges in the interactome for ribosomal and ribosome-related proteins: while these proteins account for ∼50% of both the Wt SRSF1 interactome and of the interactors lost in the FFDD mutant, they represent only ∼22% of the proteins lost in the W134A mutant. This suggests that W134A retains more interactions with proteins in this category than the FFDD mutant. Consistently, Gene Ontology analysis shows that translation-related biological processes (translation, rRNA processing, ribosome biogenesis) are significantly enriched among the proteins underrepresented in FFDD immunoprecipitates, but not in those of W134A. In contrast, protein interactors involved in mRNA metabolism–related processes are similarly altered in both mutants (Figure 3C).

To validate our quantitative SILAC proteomics analysis, we performed co-immunoprecipitation experiments using cell extracts overexpressing SRSF1-Wt or its mutants, SRSF1-W134A and SRSF1-FFDD, followed by western blot. Our results showed that U1-70K, a well-known interactor of SRSF1-Wt, continues to co-immunoprecipitate with the W134A mutant, while its interaction is reduced with the FFDD mutant (Supplemental Figure 4D). This observation aligns with our SILAC data, where U170K displayed no significant difference in binding to SRSF1-(W134A) compared to SRSF1-Wt, but exhibited decreased interaction with SRSF1-(FFDD) (vertical dashed line in Figure 3A). These findings reinforce the consistency between our SILAC results and the co-immunoprecipitation validation, even for subtle differences in interaction.

### Differential behaviour of SRSF1 RRM-mutants

The dissimilarities between the interaction landscape of each RRM-mutant could be a consequence, in part, of a differential binding of Wt and mutant SRSF1 to mRNAs leading to changes in the assembly of ribonucleoprotein complexes. RRM-mediated RNA binding ability was assigned to these specific residues by experiments with UV light-induced crosslinking of purified recombinant proteins, or domains, to pre-mRNA [11], [32]. Here, we tested this capability of FFDD and W134A mutants using RNP immunoprecipitation assays (RIP) in the context of whole cell lysates. RNA isolated from GFP IP experiments obtained from cells expressing either GFP alone, GFP-SRSF1 Wt, GFP-SRSF1 (FFDD), or GFP-SRSF1 (W134A), was used for the quantification of a variety of mRNAs by RT-qPCR, as described in the Materials and Methods. We chose to test two transcripts already described as binding targets of SRSF1, i.e., UBC9 and GAPDH [57], as well as the mRNA encoding for Akt1, the splicing of which is altered by the overexpression of SRSF1 [58], [59]. Significant enrichment of Akt1, UBC9 and GAPDH mRNAs was observed in the complexes co-purified with SRSF1 Wt, as compared with complexes co-purified with GFP alone (Figure 4A). For two of the three mRNAs (UBC9 and Akt1), the level of co-immunoprecipitation with the FFDD mutant was similar, or even higher, to that with SRSF1 Wt, while the W134A mutant showed a lower enrichment for all of the tested transcripts (Figure 4A).

**Figure 4:**
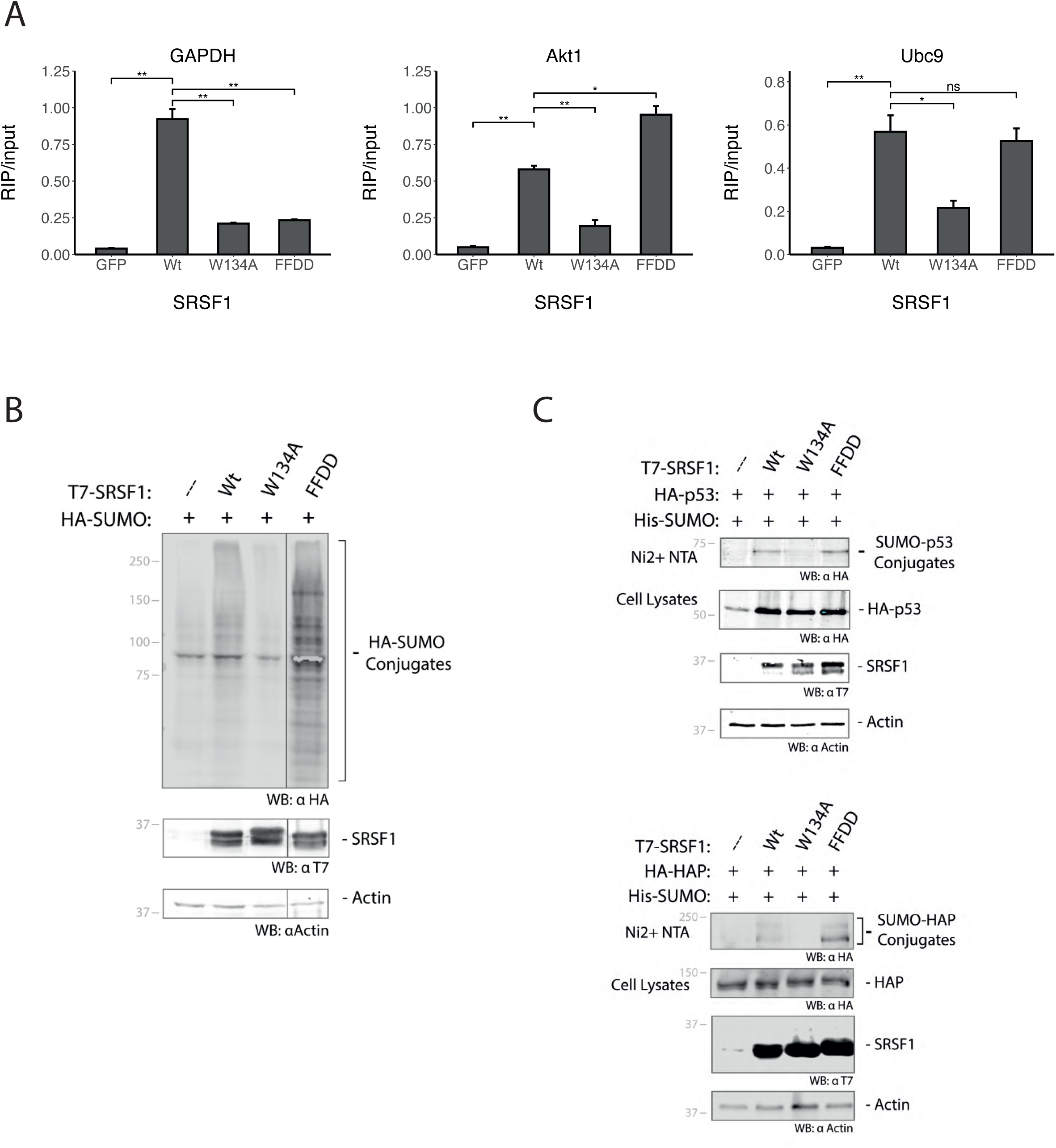
Differential behavior of SRSF1 RNA-binding mutants. (A) HEK 293T cells were transfected with expression vectors for the different SRSF1 variants fused to GFP, or for GFP alone. After 24 h, GFP-trap system was used to immunoprecipitate GFP or GFP-fusion proteins, followed by RNA extraction. After reverse transcription, GAPDH, Akt1 and Ubc9 mRNAs present in each immunoprecipitate were analyzed by qPCR, as indicated at the top of each graph. The quantification of three biological replicates is shown in a bar graph, where the brackets indicate the between-group p-value determined using a paired two-tailed t-test. Significant p-values are indicated by asterisks above the graphs (***P < 0.001; **P < 0.01; *P < 0.05). (B) Global SUMOylation with SRSF1 variants. HEK 293T cells were co-transfected with expression vectors for HA-SUMO1, and SRSF1 variants or an empty vector (---). Furthermore, an expression vector for the SUMO E2-conjugating enzyme Ubc9 was co-transfected to promote SUMOylation. Whole cell lysates were analyzed by Western blot with αHA antibodies to detect the effect of overexpressing the different SRSF1 variants on global SUMOylation patterns. (C) Analysis of the regulatory activity of the different SRSF1 variants on specific SUMO conjugation targets. HEK 293T cells were co-transfected with expression vectors for His-SUMO1, Ubc9, a known SUMOylation substrate (HA-p53 in upper panel or HA-HAP in lower panel) and different variants of T7-SRSF1 as indicated at the top of each gel. After 48 h, cell lysates were subjected to Nickel-affinity purification to enrich for SUMOylated proteins and further analyzed by Western blot with antibodies against HA. Aliquots of whole cell lysates were analyzed in order to control the expression levels of the transfected proteins.

Given that the two different SRSF1 RRM mutants showed different protein interaction networks and differential interaction with certain target mRNAs, we next tested whether they also behave differently with respect to the SRSF1 function as a positive regulator of protein post-translation modification by SUMO conjugation, a regulatory activity unrelated to its canonical role in splicing, which was previously identified [18], [60]. Thus, we tested the ability of the SRSF1 W134A and FFDD mutants to stimulate SUMO conjugation, using SRSF1 Wt as a positive control. Transient exogenous expression in HEK 293T cells of HA-tagged SUMO1 with either SRSF1 Wt, or the SRSF1 FFDD mutant, strongly stimulated global protein SUMOylation, whereas the SRSF1 W134A mutant showed no effect at all (Figure 4B). Similarly, the SUMOylation of *bonafide* SUMO conjugation substrates, including HAP/Saf-B and p53, appeared to be enhanced by over-expression of either SRSF1 Wt, or the SRSF1 FFDD mutant, but not by over-expression of the SRSF1 W134A mutant (Figure 4C). Moreover, the FFDD mutant showed a higher level of SUMOylation enhancement than SRSF1 Wt, both globally and with specific substrates analysed.

Altogether, these results indicate that the RRM-mutants, each one with a particular protein interaction network, do not always behave in the same way when challenged to different known functions of SRSF1, suggesting a differential dependence of each RRM domain for the contribution of binding partners necessary for these activities.

## DISCUSSION

SRSF1 is a well-studied auxiliary splicing factor whose multifunctional roles, extending beyond splicing, exemplify the versatility of SR proteins [44]. Its interaction with diverse partner proteins is central to these varied molecular functions. Using a SILAC-based proteomic strategy, we quantitatively distinguished specific interactors from background, generating a high-confidence interactome of 185 proteins. Approximately 27% correspond to known splicing factors and 19% to other RNA- or DNA-binding proteins (Figure 1F). Importantly, 162 of these proteins were not previously annotated as SRSF1 interactors in BioGRID, substantially expanding the current view of its interaction network (Supplemental Table 1).

Comparison with a previously reported SRSF1 interactome by the Krainer laboratory [49] revealed ∼40% overlap (Figure 1E), validating both datasets. Differences are expected due to the complementary nature of the approaches (Figure 1D) and the context-dependent multiple functions of SRSF1 across subcellular compartments. This variability is further discussed below. Despite these differences, the overall protein-category profiles are consistent (Figure 1F), and the SILAC approach appears particularly sensitive for detecting RNA-mediated interactions (Supplemental Table 1). Notably, a recent study by the same laboratory [60] employed a proximity-labeling/MS approach rather than IP-MS, and detected many interactors not previously revealed by the 2015 dataset. When we compared their list with both the 2015 findings and our own SILAC data, we likewise observed only limited overlap (10-16 proteins), further underlining that methodological variation can uncover complementary, rather than redundant, views of the SRSF1 interaction network. Together, these data underscore the multifaceted nature of SRSF1 and provide a valuable resource for future studies, including the effects of SRSF1 overexpression on protein abundance [61].

The RNA recognition motifs (RRMs), are the most common RNA-binding domains in eukaryotes [62]. A canonical domain, RRM1, and a non-canonical one, RRM2, are primarily responsible for the specific interaction of SRSF1 with RNA [32], [63]. Although structural data for RRM1 have only recently emerged [64], and pseudo-RRMs remain partially enigmatic [65], [66], their combined contribution to SRSF1 functions is still incompletely understood. To dissect the roles of each RRM, we employed full-length SRSF1 variants with point mutations F56D-F58D (RRM1) and W134A (RRM2), which preserve domain folding (Supplemental Figure 2) while disrupting RNA-binding [11] [32]. RIP assays in cultured cells confirmed loss of RNA association for W134A, whereas FFDD retained partial interaction with selected mRNAs (Figure 4A). These results could be explained in two possible ways: i) either the FFDD mutant retains its interaction with certain RNAs through other residues within RRM1, or other parts of the protein, or, ii) it continues to interact with several proteins in stable ribonucleoprotein complexes retaining an indirect interaction with the tested mRNA. The marked loss of mutant FFDD protein-protein interactions discussed below (Figure 3A and Supplemental Figure 4B), makes us favour the first scenario.

Since RNA binding is tightly coupled to SRSF1’s role in coordinating macromolecular complexes, we next examined how disrupting each RRM affects its network of protein interactions. Consistent with prior studies [32], GFP-tagged mutant constructs were transiently overexpressed without silencing endogenous Wt SRSF1 to preserve essential cellular functions [44], and minimize indirect effects on global gene expression [9]. Only exogenous proteins were immunoprecipitated, so endogenous SRSF1 may compete for binding partners, which could reduce the apparent number of mutant-specific interactions, but it provides a more physiological context for comparing the association landscapes of Wt and mutant SRSF1 proteins. SILAC analyses revealed distinct association profiles: W134A caused moderate loss of interactions, while FFDD led to widespread reduction (Figure 3A). This is in agreement with observations for SR34, an SRSF1 ortholog in *Arabidopsis*, where an RNP1 mutant, designed with the same criteria as the FFDD, loses interactions that are preserved with an RRM2 mutant designed with the same criteria as W134A [67]. Although it has already been widely proposed that RRM2 is an important mediator of protein-protein interaction [14], [68], our results highlight the central role of RRM1 in shaping the SRSF1 interactome, consistent with structural and phosphorylation-dependent regulatory effects on RRM1 surfaces [69].

The loss of interactions may reflect both RNA-dependent and RNA-independent mechanisms. The nuclease-sensitive interactions detected by Akerman *et al*. [49] could be an example of the first point. Regarding the second, many associations not mediated by RNA have been reported [15], [16]. An example of this RNA-independent protein binding, could be the interaction of SRSF1 with the SR protein kinase 1 (SRPK1), with which residues W134 and Q135 make direct contact [70]. However, for this particular case we have not detected significant differences between SRPK1 interaction with W134A or with Wt SRSF1, suggesting that other residues may also contribute to stabilize SRSF1 binding to this kinase, similar to what was previously reported by Nhat Huynh *et al*. [71]. Curiously, SRPK1 was significantly less immunoprecipitated with the FFDD variant (Figure 3A), emphasizing anew the importance of RRM1 for protein interactions. Whether in the case of a direct, indirect, or RNA-dependent binding, the loss of interaction with certain proteins will undoubtedly shape the role of the SRSF1 variant in the activity under study.

The global analysis of the molecular processes in which these less-interacting proteins are involved, also showed differences between SRSF1 (FFDD) and (W134A). The results shown in Figure 3B and 3C, when compared with Figure 1F, suggest that RRM1 mutant loses selectively more interactions with translation-related proteins than with DNA/RNA regulators, whereas RRM2 mutant seems not to specifically lose the interactors related to translation. These interesting observations not only reinforced a strong difference between both mutants, but also reveal a clear distinction in the contribution of each RRM domain to the macromolecular complexes in which SRSF1 takes part, with a tendency to suggest that RRM1 is curiously more responsible for translation-related activities of SRSF1 than RRM2. It is worth clarifying that, despite these broad differences, the identity of the proteins that interact less with each particular mutant could independently affect some mechanisms of these categories. For instance, the role of SRSF1 in the activation of translation initiation by 4E-BP regulation has been shown to depend on its RRM2 [15]. Even more, a mutant in which the WQD residues in the conserved heptapeptide have been replaced by three alanines (SRSF1-AAA), does not interact with components of the mTOR pathway, failing to enhance translation [14], [15].

The few experiments published to date in which both mutants have been used simultaneously did not show differences between them. In addition to the fact that these RNA-binding mutants do not show differences between them in their localization or in the prototypic splicing activity (Figure 2 and [35]), neither do so for the ESE binding [34]. Furthermore, SRSF1-AAA mutant (related to W134A) behaves similarly to FFDD in the loss of ability to promote β-catenin accumulation [40]. Nevertheless, the different association landscape of each mutant lead us to speculate that they could not always behave in the same way. Evidence of this, is the here reported opposite effect of both variants in the regulation of SUMO conjugation (Figure 4B-C). Various studies have been performed with the FFDD, W134A and AAA variants, several times using them just as generic RNA-binding mutants. Many of the results from those studies may even seem contradictory. An example is the reported *in vitro* proper functioning of SRSF1 (FFDD) in alternative splicing [32], opposite to what we detected here with the transfection of the reporter minigene (Figure 2C). The complexity of these factors, together with the still unclear molecular mechanisms underlying many of these activities as well as the technical differences of each experimental setting, could explain some of the apparent discrepancies in the literature, and highlight the importance of contextual and mechanistic dissection of SRSF1 functions using domain-specific variants. Based on what we report here, complementing previously available bibliographic data, future use of these mutants will have to take into account not only the possibility of this differential behaviour, but also the fact that each of them loses interactions with a distinct subset of proteins, either directly or indirectly. These differential interactomes suggest that the functional consequences of RNA-binding impairment may go beyond the mere loss of RNA recognition and depend on specific protein partnerships affected in each case. Further functional studies with both mutants simultaneously will be required to elucidate the contribution of each RRM to the diverse activities of SRSF1.

Comparison of the two SILAC experiments revealed incomplete overlap among Wt interactors, which is expected because each constitutes an independent analysis with distinct biological samples and MS runs. SILAC is optimized for within-experiment comparisons rather than generating exhaustive interactor lists, so proteins below the threshold in SILAC 1, when comparing Wt to GFP, may be significant in SILAC 2 when comparing Wt to mutants. Technical variability and sensitivity to experimental conditions further contribute to these differences. These complementary datasets, summarized together in Figure 5, provide a more comprehensive view of the SRSF1 interactome, capturing distinct dimensions of RNA-binding-dependent and independent interactions.

**Figure 5:**
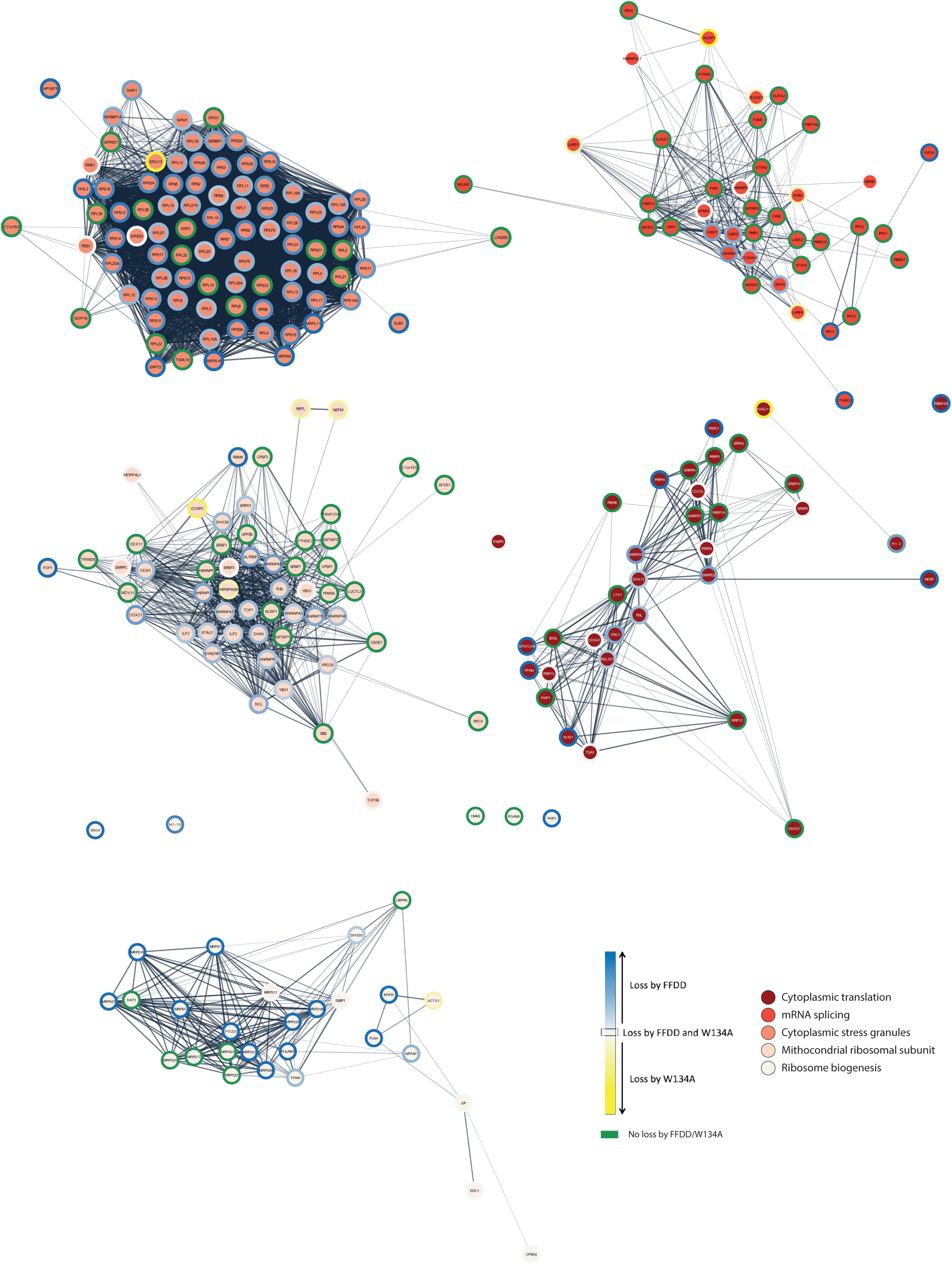
Protein-protein association landscape of SRSF1 and its RNA-binding mutants. STRING-based interaction network of SRSF1 interactors and proteins that show reduced association with SRSF1 RNA-binding mutants W134A and FFDD, compared to wild-type (Wt) SRSF1. Five functional clusters, defined by STRING clustering, are highlighted by node fill colors. Each node represents a protein and is outlined according to its interaction loss profile: blue contours indicate decreased binding to W134A (with colour intensity reflecting interaction loss magnitude), yellow contours indicate reduced binding to FFDD, white contours denote proteins with diminished association to both mutants, and green contours indicate an SRSF1 partner that doesn’t lose interaction with any of the RNA-binding mutants.

The intricate behaviour of RRMs, capable of interacting with RNA, DNA, and other proteins [26], renders the dissection of SRSF1 functions challenging. The present dataset enables construction of an atlas of protein interactions for Wt and RRM-mutants SRSF1 (Figure 5), facilitating future analyses of macromolecular complexes and their dependence on individual RRMs. This may represent extremely useful information to disentangle the molecular mechanisms underlying the many functions SRSF1 is involved in. Beyond SRSF1 itself, this work provides insights into RRM domain behaviour, which is of broad interest due to the regulatory versatility of these motifs and their potential as therapeutic targets [72]. Taken together, our data redefine the functional landscape of SRSF1 by revealing domain-specific interaction networks that underlie its pleiotropic roles.

## ACKNOWLEDGEMENTS

The authors thank Valeria Buggiano, José Clemente and Amaranta Avendaño for technical assistance, as well as the members of the Petrillo, de la Mata, Muñoz, Schor and Kornblihtt laboratories for their stimulating interactions. We are grateful to Graciela Boccaccio, Julio Caramelo and Federico Pelisch for their valuable suggestions, and to Alexandra Elbakyan for the support of Latin American science. We especially thank Guillermo Risso for his inspiring ideas and encouraging discussions.

## MATERIALS AND METHODS

### Cell lines

HeLa (human cervix adenocarcinoma cell line, ATCC CCL-2) and HEK 293T cells (human embryonic kidney cell lines, ATCC #CRL-11268) were maintained in Dulbecco’s modified Eagle’s medium. All the media were supplemented with 10% fetal bovine serum, 100 U/ml penicillin, and 100 μg/ml streptomycin, and the HEK 293T medium was also supplemented with 110 mg/L sodium pyruvate.

### DNA plasmids and transfection

The plasmid used were T7-SRSF1 (provided by A. Krainer and J. Caceres), HA-SUMO1 and His-SUMO2 (provided by R. Hay), V5-UBC9 (provided by E. Artz), HA-HAP/Saf-B (provided by G. Biamonti), HA-p53, GFP-SRSF1 Wt (provided by A. Lamond), pcDNA 3.1 (+) (Addgene #2093), pEGFP-C1 (Addgene #2487) and EDI minigen (provided by A. R. Kornblihtt).

T7-SRSF1 W134A, T7-SRSF1 FFDD, GFP-SRSF1 W134A, GFP-SRSF1 FFDD were generated in our laboratory by site-directed mutagenesis from the respective Wt backbones. This was performed by the DpnI method, based on Stratagene’s QuickChange specifications. Both the expected mutations and the absence of off-target mutations were always verified by sequencing.

Plasmid DNA was transfected into HEK 293T or HeLa cells with Lipofectamine 2000 according to manufacturer’s instructions (Invitrogen).

### Stable isotope labeling by amino acids in cell culture (SILAC)

#### Generation of labelled cell lines

DMEM media without lysines and without arginines were supplemented with 10% serum, penicillin/streptomycin, and separated into two or three groups. The amino acids with differential isotopes (Sigma) were added to each one as follows: arginine R0 and lysine K0 for light cells, arginine R6 and lysine K4 for medium cells, and arginine R10 and lysine K8 for heavy cells.

An initial culture of HEK 293T cells was spread into three 10cm dishes. Each of them was incubated with one of the media (light, medium, heavy). Six passages of each line were carried out, always maintaining the media with the corresponding isotopes, in order to obtain a total protein labelling in each group. In each passage, the cells were lifted up-down with a pipette, and trypsin was not used since it may have unlabeled amino acid residues. 24 hours before transfection, 3.5×10^6^ cells for each experimental condition were plated in 10 cm plates.

#### Transfection and harvest of cells with GFP-SRSF1 constructs

HEK 293T cells labeled with amino acid isotopes were transfected with Lipofectamine 2000 (Invotrogen), according to the protocol mentioned above. For each condition, 12 μg of the expression plasmid for the corresponding GFP-SRSF1 version, or GFP alone, were used. After the addition of the complexes, the cells were left at 37°C and 10% CO_2_ for 24 hours.

Cells were harvested by up-down semi-automatic pipetting in their own culture media. The cells were then quantified, and the same amount was taken in each condition. These cells were centrifuged at 500g for 5 minutes at 4°C, and finally washed twice with cold PBS.

#### Immunoprecipitation and sample preparation

The iST GFP-Trap kit (ChromoTek, PreOmics) was used, following the protocol provided by the manufacturer.

#### Liquid chromatography (LC)-tandem mass spectrometry (MS/MS)

Prior to MS-based analyses, all peptide samples were desalted on self-made Empore C18 (Agilent Technologies, cat. no. 12145004) Stop and Go Extraction Tips (StageTips) according to a protocol by Rappsilber et al. (2007). Desalted peptide samples were analysed using EASY-nLC 1000 nanoflow UHPLC system, EASY-Spray ion source and Q Exactive hybrid quadrupole-Orbitrap mass spectrometer (all Thermo Fisher Scientific). Peptides were loaded onto 2 cm Acclaim PepMap 100 C18 nanoViper pre-column (75 mm ID; 3 mm particles; 100 A_°_ pore size) at a constant pressure of 800 bar and separated using 50 cm EASY-Spray PepMap RSLC C18 analytical column (75 mm ID; 2 mm particles; 100 A_°_ pore size) maintained at 45_°_C.

Exploratory analysis using standard MS settings was performed using 10% of the sample and peptides were separated with 60 min linear gradient of 5–22% (vol/vol) acetonitrile in 0.1% (vol/vol) formic acid at a flow rate of 250 nl/min, followed by a 12 min linear increase of acetonitrile to 40% (vol/vol). Total length of the gradient including column washout and re-equilibration was 90 min. Peptides corresponding to the complete human proteome were analysed with 240 min linear gradient of 5–22% (vol/vol) acetonitrile in 0.1% (vol/vol) formic acid at a flow rate of 250 nl/min.

Peptides eluting from the liquid chromatography column were charged using electrospray ionisation and MS data were acquired online in a profile spectrum data format. Full MS scan covered a mass range of mass-to-charge ratio (m/z) either 300–1800, or 300–1600, during standard and comprehensive peptide analyses, respectively. Target value was set to 1,000,000 ions with a maximum injection time (IT) of 20 ms and full MS was acquired at a mass resolution of 70,000 at m/z 200. Data dependent MS/MS scan was initiated if the intensity of a mass peak reached a minimum of 20,000 ions. During standard LC-MS/MS analyses, up to 10 most abundant ions (Top 10) were selected using 2 Th mass isolation range when centered at the parent ion of interest. For comprehensive analyses, up to 4 (Top 4) most abundant ions were picked for MS/MS. Selection of molecules with peptide-like isotopic distribution was preferred. Target value for MS/MS scan was set to 500,000 ions with a maximum IT of 60 ms and resolution of 17,500 at either m/z 200 for standard, or maximum IT of 500 ms and a resolution 35,000 at m/z 200 for comprehensive peptide analyses. Precursor ions were fragmented by higher energy collisional dissociation (HCD), using normalised collision energy of 30 and fixed first mass was set to m/z 100. Precursor ions with either undetermined, single, or high (>8) charge state were rejected. Ions triggering a data-dependent MS/MS scan were placed on the dynamic exclusion list for either 40 s (standard analyses), or 60 s (comprehensive analyses) and isotope exclusion was enabled. The sample was analyzed twice according to the settings of the comprehensive analysis.

#### Data analysis

To detect the significantly enriched proteins in the immunoprecipitate of each condition, data analysis was performed according to Emmott E. *et al.* [73] and Have S*. et al*. [74]. Graphs, regressions and statistics were made in RStudio.

### KEGG pathway enrichment and visualization

KEGG pathway enrichment analysis was performed using R (v4.5) within the RStudio environment. The analysis employed the clusterProfiler, pathview, and org.Hs.eg.db packages from Bioconductor [75], [76]. Protein identifiers were converted to Entrez Gene IDs and used to identify enriched pathways from the KEGG database using an adjusted p-value cutoff of 0.05 (Benjamini-Hochberg correction). Visualization of enriched pathways, including pathway maps such as the Ribosome pathway, was carried out using the pathview function with default settings.

### Gene Ontology analysis

For the gene ontology (GO) analysis, the DAVID (Database for Annotation, Visualization and Integrated Discovery) bioinformatics platform was used [77], [78]. This analysis was done with respect to three classes: biological processes (BP), cellular component (CC) and molecular functions (MF). For each category within these classes, the fold enrichment of the sample relative to the number of genes expected in a random selection was calculated. For each enrichment, the significance of the difference was studied through the p-value parameter.

### Structural modeling and visualization

Predicted structures of the RNA recognition motifs (RRMs) of SRSF1 were obtained using AlphaFold 3 (AlphaFold Server) [79]. The sequences corresponding to the RRM1 and RRM2 domains (aa 11–91 for RRM1, 121–195 for RRM2) were used as input. Mutant versions were generated by introducing the desired amino acid substitutions (F56D, F58D for RRM1; W134A for RRM2) into the input sequence. Resulting models were analyzed and visualized with UCSF ChimeraX (version 1.10.1) [80]. Wild-type and mutant structures were superimposed using MatchMaker to evaluate potential structural deviations. Residues of interest were highlighted and color-coded (green for wild-type, yellow for mutant). Structural elements such as α-helices and β-sheets were represented using ribbon diagrams, with additional stick representations for side chains at mutated positions.

### SDS-Page and silver stain

Cells transfected as described were harvested, washed with PBS and lysed in 1× Laemmli sample buffer. Protein samples were resolved using a 4-12% NuPAGE Bis-Tris gel (Life Technologies). The gel was fixed for 20 minutes in 150 ml of fixation buffer (50% methanol, 5% acetic acid), followed by two 10 min washes, once in 50% methanol and once in water. For the sensitization the gel was incubated 1 minute with 150 ml of 0.02% Sodium Thiosulfate, and then rinsed twice with water. The gel was immersed in 150 ml of 0.1% silver nitrate with 0.08% formalin (37%) for 20 minutes. Development was performed by incubation in a fresh 2% sodium carbonate solution with 0.04% formalin (37%) until the desired staining intensity. Staining was stopped by washing the gel in 5% acetic acid for 10 minutes, and finally washed with water.

### Fluorescence Microscopy and imaging

HeLa cells were plated in multiwell culture dishes over glass microscope coverslips. The next day, cells were transfected with expression plasmids for SRSF1 variants fused to GFP. At 24 hours, the cells were fixed in 3.7% paraformaldehyde in 37°C PHEM buffer (60 mM Pipes, 25 mM Hepes, 10 mM EGTA, and 2 mM MgCl_2_, pH 6.9) for 5 minutes. Subsequently, they were permeabilized with 1% Triton X-100 in PBS for 10 minutes. For nuclei staining, the TOTO-3 dye was used for 20 minutes at room temperature, previously treating the cells with 0.1 mg/μl RNase at 37°C for half an hour. Finally, the cells were washed with PBS and mounted in VECTASHIELD medium (Vector Labs). Images were acquired with an Olympus FV300 confocal microscope, with a 60X 1.4 NA objective.

To quantify the fluorescence signal from a large number of microscopy images, CellProfiler software (v3.1.5) was used. A command line was developed to automatically recognize the nuclei and speckles observed in each photograph. To achieve this, the appreciable difference in intensities between both compartments was used. Establishing a minimum and maximum diameter for each object (nucleus or speckle), an intensity limit was assigned to differentiate each structure as an independent object. Next, a relationship was established to associate each speckle with a certain nucleus. Once these were recognized, the “nucleoplasm” object was identified as the subtraction of the speckles to the associated nucleus. The quantification was performed by analysing the ratio between the mean intensities of the speckles of a certain nucleus, and that of the corresponding nucleoplasm (obtained by subtracting the speckles from the nucleus).

### Fluorescence recovery after photobleaching (FRAP)

HeLa cells were plated in multiwell culture dishes over microscopy glass coverslips. The next day, cells were transfected with expression plasmids for SRSF1 variants fused to GFP. 24 hours later, cells were imaged on an Olympus U-Plan-Apo microscope with a 100×1.35NA objective and photobleached using the DeltaVision Spectris microscope’s photokinetic experiment function. A small region within the nucleus was photobleached with a 488nm laser (100% power for 0.15 seconds), and photographs were taken at 300mSec intervals. The intensities of the selected areas, before and after photobleaching, were quantified using ImageJ software.

The fluorescence signal was quantify as indicated above.

For data analysis, we proceeded as follows with the images of each speckle: each intensity measurement on the area of interest was relativized to a point outside this area (to avoid the effects of change of focus between each image, photobleaching by exposure, and decrease in the population of fluorescent proteins by photobleaching); an average of the pre-photobleaching intensities was made in the area of interest; each post-photobleaching point was normalized to the average number of pre-photobleaching points; these values were plotted as a function of post-photobleaching time; the data was fitted to the following exponential function that describes the recovery dynamics [*fluorescence intensity = k_1_(1-ê(-time/k_2_))+k_3_*], where *k_1_* represent the mobile fraction and *k_2_* the diffusion coefficient. The graphs of the recovery curves, and the different graphs of the adjustment parameters, as well as the statistical analyzes were carried out with RStudio.

### RNA extraction, RT, end point- and real time-PCR

Total cellular RNA was isolated by using Tri-Reagent (Invitrogen) and reverse transcribed with random deca-oligonucleotide primer mix. End-point PCR was performed with dCTP alpha-32P. Radioactive PCR products were run on native 6% polyacrylamide gels, which were subsequently dried and exposed to X-ray films (Agfa). Radioactivity in the corresponding bands was measure by scintillation counter and used to calculate the radio between different mRNA isoforms.

Quantitative PCRs (qPCRs) were performed by using SYBR Green dye and specific primers for different mRNAs. The specific primers used were: TTGTGAAGGAGGGTTGGCTG (hAkt1 – FW), CTCACGTTGGTCCACATCCT (hAkt1 – REV), CAATGACCCCTTCATTGACC (GAPDH – FW), GATCTCGCTCCTGGAAGATG (GAPDH – REV), AAGTCCCGAGACAAAGGGAAG (Ubc9 – FW), GGTGACTAGTCATTGTATGGAG (Ubc9 – REV).

### Western blots

Protein samples were resolved by SDS-PAGE and transferred to nitrocellulose membranes (GE Healthcare Amersham). Membranes were blocked for 1 hour with a 5% milk in TBS buffer, and then incubated with primary antibodies. After washing, membranes were incubated with IRDye® 800CW or 680RD (LI-COR Biosciences) secondary antibodies. Bound antibody was detected using an Odyssey imaging system (LI-COR Biosciences). Western blots were performed at least three times from independent experiments and representative images are shown in each case. The antibodies used were: mouse monoclonal anti-β-actin C4 (Santa Cruz Biotechnology, sc-47778), mouse monoclonal anti-HA.11 (Covance, MMS-101P), mouse monoclonal anti-T7 (Novagen, 69522), mouse monoclonal anti-GFP B-2 (Santa Cruz, sc-9996), rabbit polyclonal anti-U1A-70K (Abcam ab83306).

### Immunoprecipitation of RNP complexes (RIP)

HEK 293T cells (3×10^6^) were harvested in their own culture media 24-36 hours after being transfected. After centrifugation at 500g for 5 minutes at 4°C, they were washed twice with cold PBS. The cell pellet was resuspended in 500 μl of IP buffer (Tris HCl pH 7.5 50mM, NaCl 150mM, Nonidet P-40 1%, Deoxycholate Na 0.5%) supplemented with protease inhibitors (Complete, Roche) and phosphatase inhibitors (5mM β-glycerophosphate and 5mM KF). Subsequently, the samples were sonicated on ice by three pulses of 20% amplitude. After an incubation of 30 minutes at 4°C, the samples were centrifuged at 4000g for 20 minutes at 4°C. 10% of the supernatant volume was taken as input, and the rest (cellular lysates) were used for immunoprecipitation assays. From 10% input, half of the volume was analyzed by western blot with anti-GFP antibody and the rest subjected to RNA extraction and RT-qPCR.

Cell lysates were incubated for 1 hour at 4°C in rotation with 10 μl of GPF-trap®, previously equilibrated with IP buffer. The microspheres were subsequently washed four times with Wash buffer (Tris HCl 7.5 50mM, NaCl 250mM, Deoxycholate Na 0.5%, NP-40 0.1%) supplemented with phosphatase inhibitors. In the last wash, 5% of the volume was taken for protein immunoprecipitation control by Western Blot. The remaining beads were resuspended in 250 μl of Tri-Reagent (Invitrogen) for subsequent RNA extraction and RT-qPCR.

### Purification of 6xHis-SUMO-conjugated proteins

HEK 293T or HeLa cells were transfected in 35 mm culture wells with the indicated plasmids. After 48 h, 6xHis-SUMO conjugates were purified under denaturing conditions using Ni-NTA agarose beads according to the manufacturer’s instructions (QIAGEN). Transfected cells were harvested in ice-cold PBS plus 100 mM iodoacetamide. An aliquot was taken as input, and the remaining cells were lysed in 6 M guanidinium-HCl containing 100 mM Na_2_HPO_4_/NaH_2_PO_4_, 10 mM Tris-HCl pH 8.0, 5 mM imidazole, and 10 mM iodoacetamide. Samples were sonicated to reduce the viscosity, and after centrifugation for 20 min at 12 000 g, proteins in the supernatants were purified using Ni-NTA beads (QIAGEN) according to Tatham *et al*. [81] Samples were subsequently washed with wash buffer I (8 M urea, 10 mM Tris-HCl, 100 mM Na_2_HPO_4_/NaH_2_PO_4_, 5 mM imidazole, 10 mM iodoacetamide, pH 8), wash buffer II (8 M urea, 10 mM Tris-HCl, 100 mM Na_2_HPO_4_/NaH_2_PO_4_, 0.2% Triton X-100, 5 mM imidazole, 10 mM iodoacetamide, pH 6.3), and wash buffer III (8 M urea, 10 mM Tris-HCl, 100 mM Na_2_HPO_4_/NaH_2_PO_4_, 0.1% Triton X-100, 5 mM imidazole, 10 mM iodoacetamide, pH 6.3). Samples were eluted in 2× Laemmli sample buffer containing 300 mM imidazole during 5 min at 95 °C.

### Statistical analysis and data visualization

All statistical analyses and data visualizations were performed in R (version 4.5) using the RStudio integrated development environment. Unless otherwise stated, standard functions from base R or packages from the Bioconductor and CRAN repositories were used. Graphs were generated using the ggplot2, ggpubr, rstatix, and minpack.lm packages.

### Protein-protein interaction network analysis

Protein interaction networks were built using Cytoscape [82]. Protein identifiers from the mass spectrometry analysis were input into the STRING app to retrieve high-confidence interaction data (confidence score ≥ 0.4) from the STRING database [83]. Functional clustering of proteins within the network was performed using the ClusterMaker app with the Markov Clustering (MCL) algorithm [84]. Network visualization and annotation were refined using the Omics Visualizer app to overlay quantitative proteomics data [85]. Node attributes (e.g., size and color) were mapped according to enrichment or statistical significance to enhance interpretability.

