## Supplemental for "Differential contribution of the RNA recognition motifs to SRSF1 protein interaction network"

#### SUPPLEMENTAL FIGURE LEGENDS

**Supplemental Figure 1: KEGG pathway enrichment analyses of SRSF1-associated proteins.** (A) Enriched KEGG pathways identified from the list of SRSF1-interacting proteins obtained in our proteomic analysis using the STRING database. Pathways related to RNA processing (e.g., spliceosome, mRNA surveillance) are significantly overrepresented. The red color indicates the protein, or protein group, identified in the analysis. (B) KEGG Ribosome pathway map highlighting the proteins detected in both our SRSF1 interactome and in the proteomic dataset from Akerman *et al*. (2015). Shared proteins are marked in red, illustrating the robust association of SRSF1 with components of the translational machinery across independent studies.

**Supplemental Figure 2: Predicted structural models of the RNA recognition motifs (RRMs) of SRSF1 and their corresponding point mutants.**

AlphaFold-based predictions are shown for RRM1 (upper panel) and RRM2 (lower panel). Structures are represented as ribbon and stick diagrams, evidencing the α-helices and β-sheets characteristic of RNA-binding domains. In each case, the wild-type (Wt) domain is displayed in white and the mutant domain in red, with both models superimposed to highlight the absence of detectable structural differences between variants. Residues subjected to mutagenesis are highlighted: phenylalanines 56 and 58 (F56, F58) in RRM1 and tryptophan 134 (W134) in RRM2 are depicted in green, while their mutant counterparts (aspartic acids in RRM1 and alanine in RRM2) are shown in yellow.

**Supplemental Figure 3: Effects of SRSF1 mutants on mobility dynamics within nuclear speckles and on alternative splicing.**

(A) Box plots displaying the parameters of mobile fraction (left panel) and diffusion coefficient (right panel) for all speckles analyzed by FRAP with each version of SRSF1. The cross line inside the boxes represents the median of each sample. The asterisks above each box show significant differences with respect to the Wt for a t-test. (****) p-value<0.0001. (B) HeLa cells were co-transfected with the indicated mini-gene and the expression vectors for different SRSF1 variants, or an empty pcDNA. The expression levels of each of these proteins were analyzed by western-blot assays. (C) RT-end point PCR analysis of splice variants derived from the fibronectin splicing reporter mini-gene as described in Figure 3C. Two representative autoradiographs for each SRSF1 variant and pcDNA control are shown. All of them are different lanes from the same gel.

**Supplemental Figure 4: Interactomes of SRSF1 RNA-binding mutants.**

HEK 293T cells were transfected with expression vectors for GFP-SRSF1 Wt, GFP-SRSF1 W134A or GFP-SRSF1 FFDD. Cell lysates were subjected to immunoprecipitation with GFP-trap. (A) Schematic representation of the experimental design for the SILAC assay. A unique cell culture was split into three, each incubated with distinct isotopically labeled amino acids over several passages to ensure complete incorporation. Cells were then transfected with expression plasmids for GFP-SRSF1 Wt,

GFP-SRSF1 W134A or GFP-SRF1 FFDD, processed, and lysed. GFP immunoprecipitation was performed on the lysates, and the resulting immunoprecipitates were combined. Finally, samples were analyzed by mass spectrometry to quantify the ratio of peptides labeled with one isotope relative to those labeled with the other. (B) Comparison between a representative replicate of immunoprecipitates for each variant, monitored by SDS-page and subsequent silver staining. (C) GO analysis of proteins with which mutants lose interaction compared to SRSF1 Wt. The diameter of the circles represents the enrichment factor of these groups, and the color indicates the p-value of the statistical analysis, according to the legend on the right panel. (MF) Molecular Functions. (D) Cell lysates expressing GFP, GFP-SRSF1 Wt, GFP-SRSF1 W134A or GFP-SRSF1 FFDD were subjected to immunoprecipitation with GFP-trap. With the immunoprecipitates and an aliquot of the pre-IP lysates (Input) a western blot was performed revealing the membrane with the antibodies indicated below each image.

**Supplemental Table 1:** SILAC-derived enrichment ratios for proteins co-precipitating with SRSF1 WT versus GFP control.

### Supp Figure 1

A

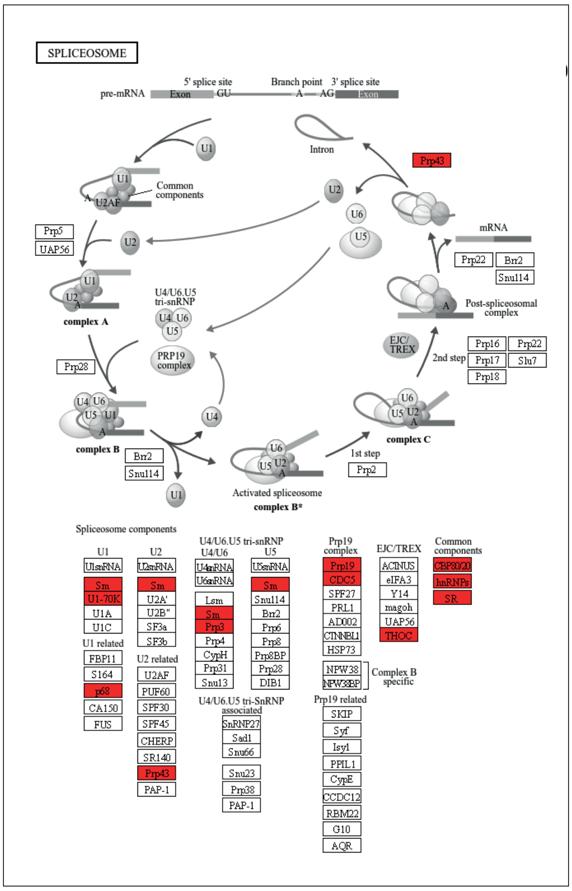


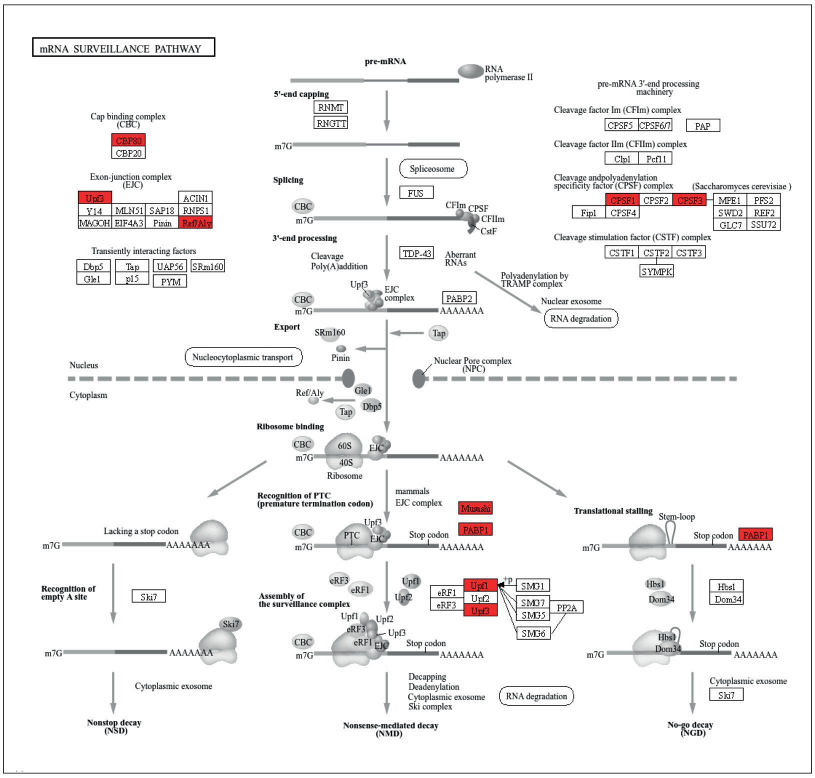


B

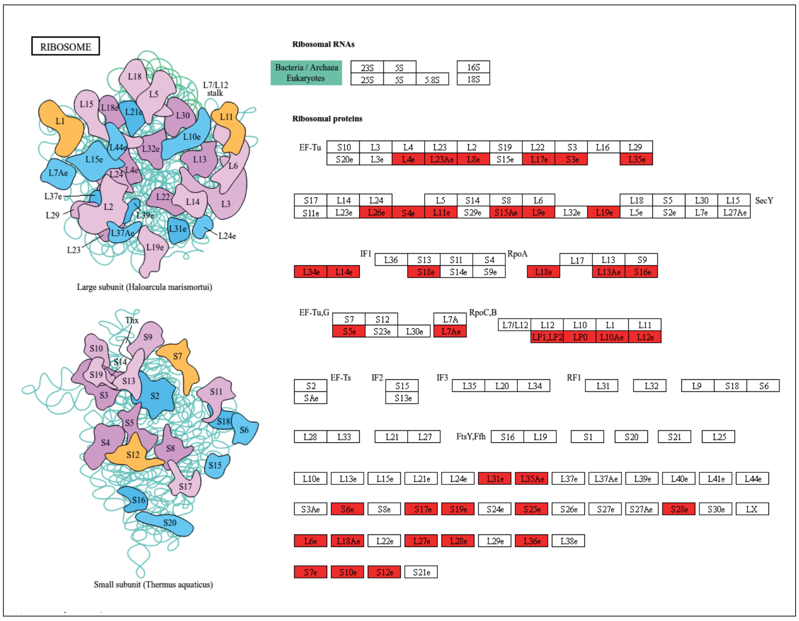

Supp Figure 2

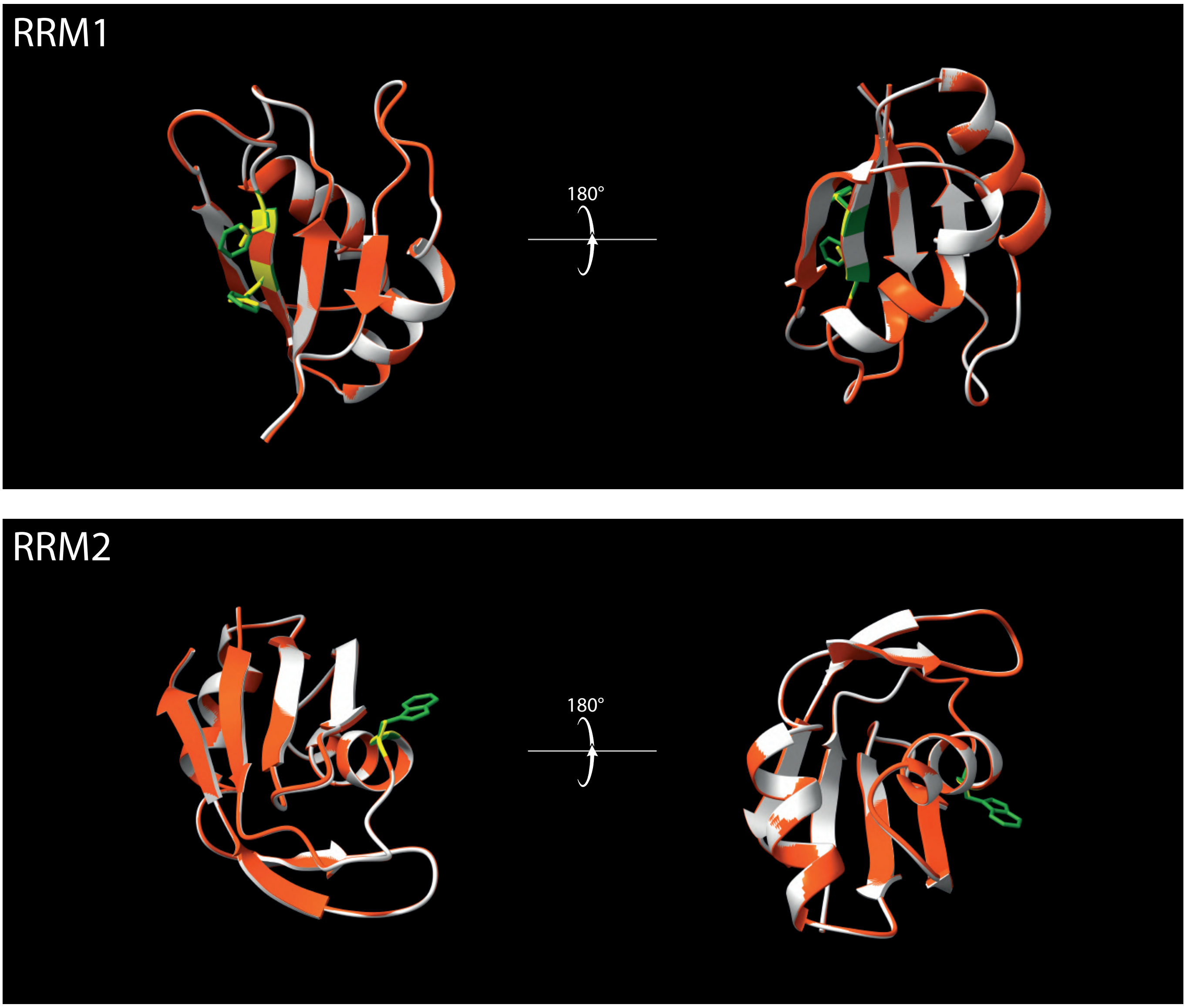

### Supp Figure 3

A

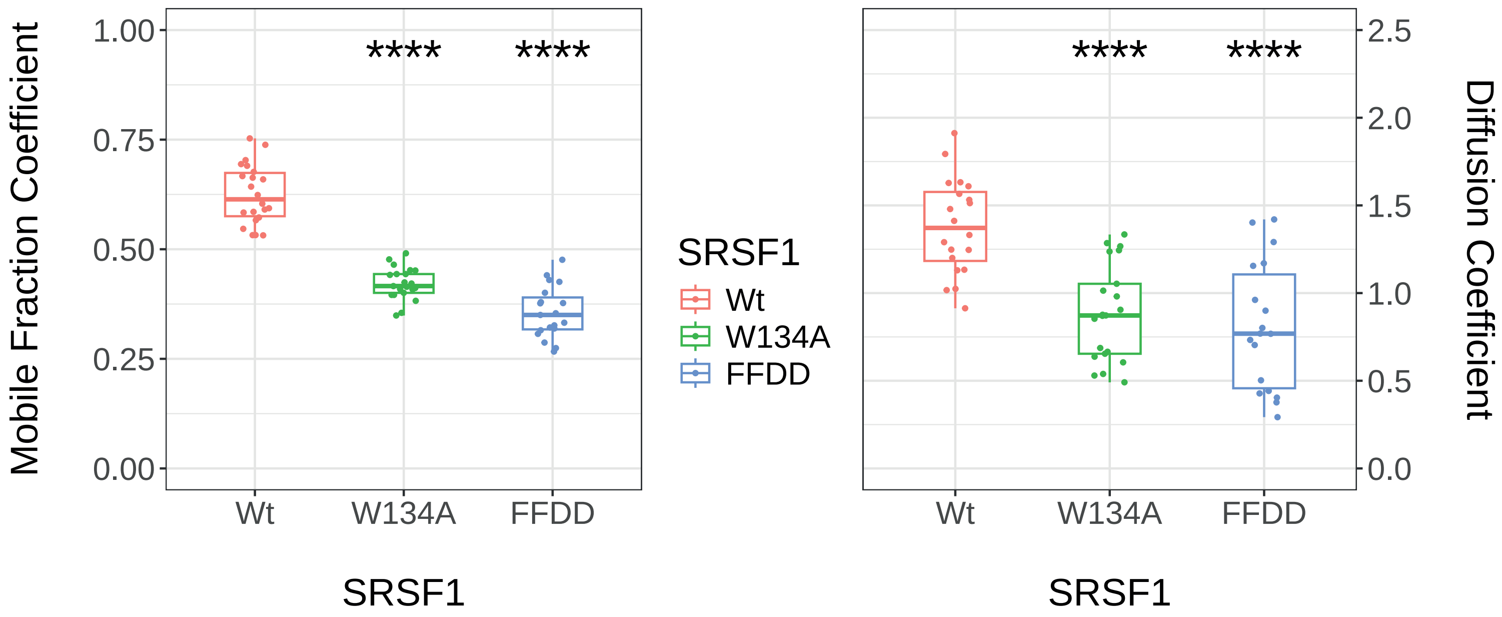


B

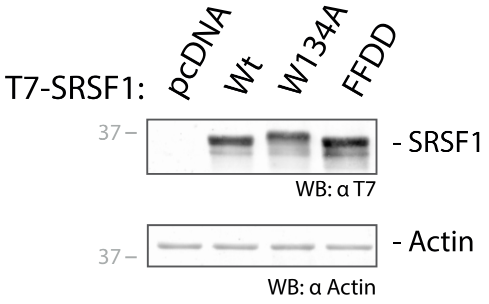


C

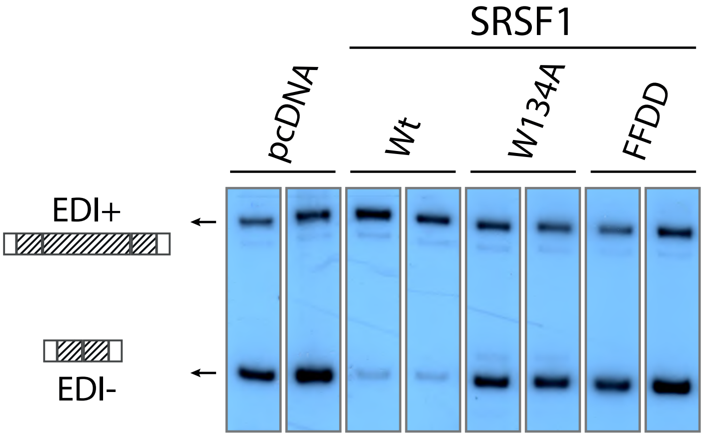

### Supp Figure 4

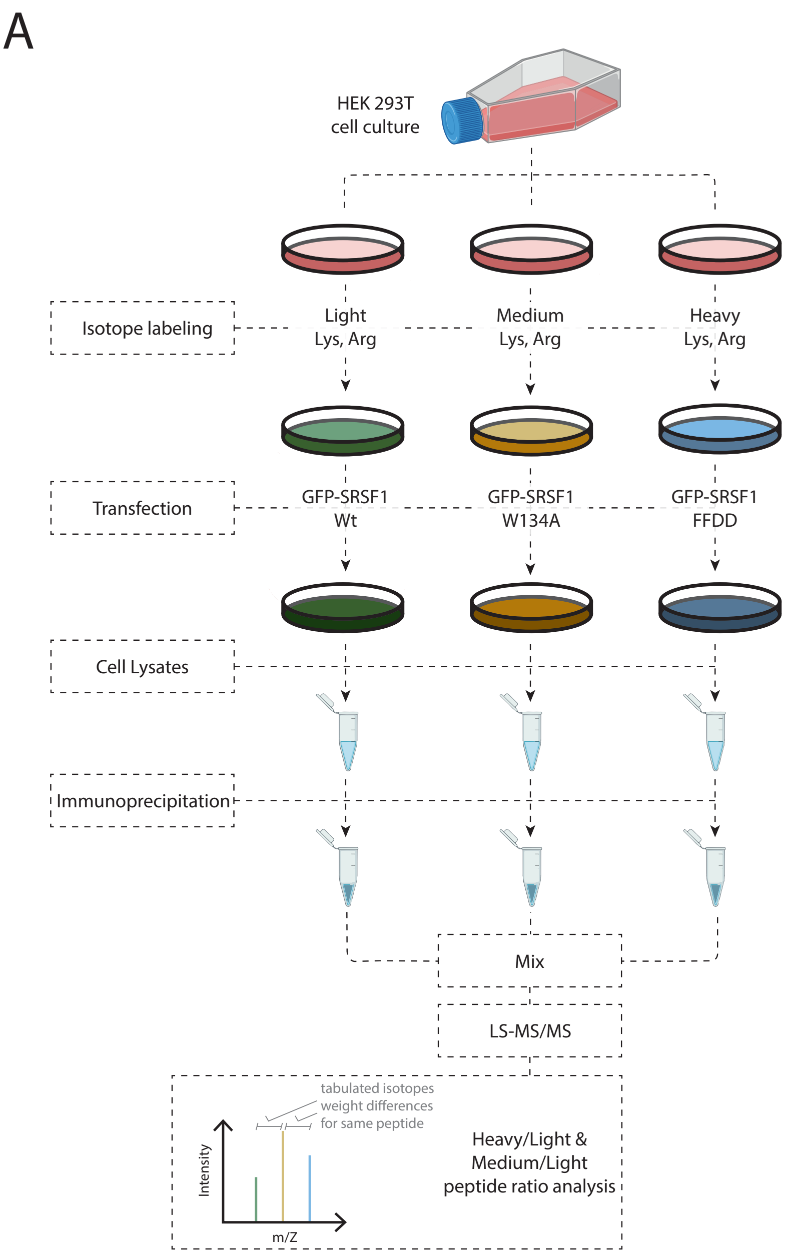


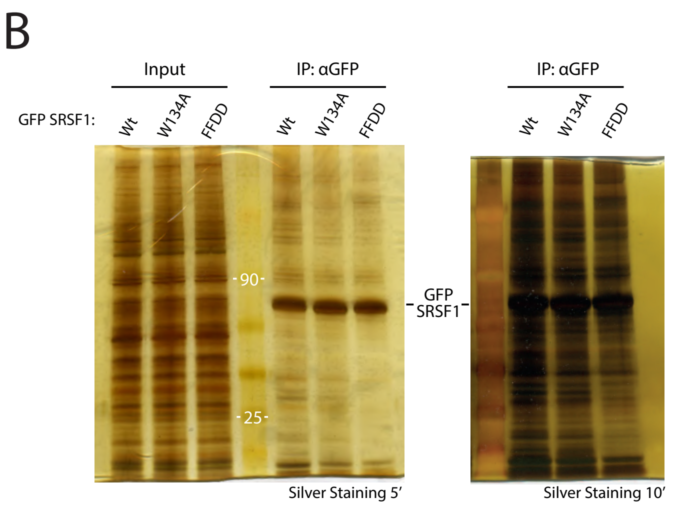


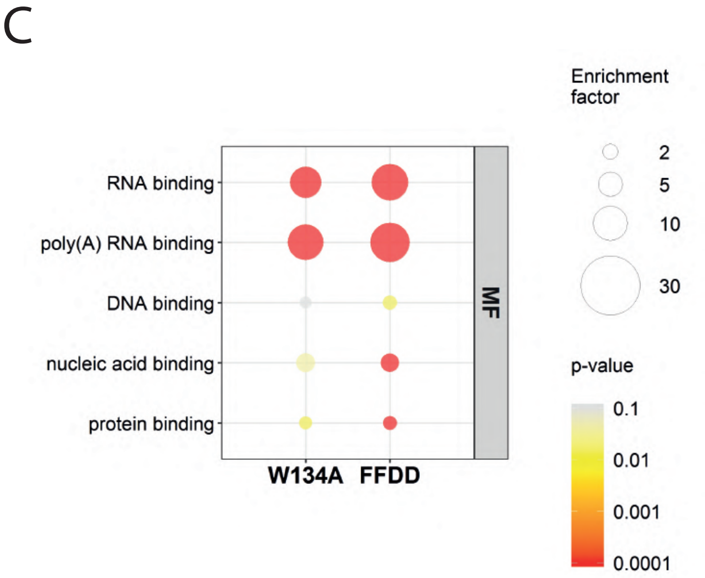


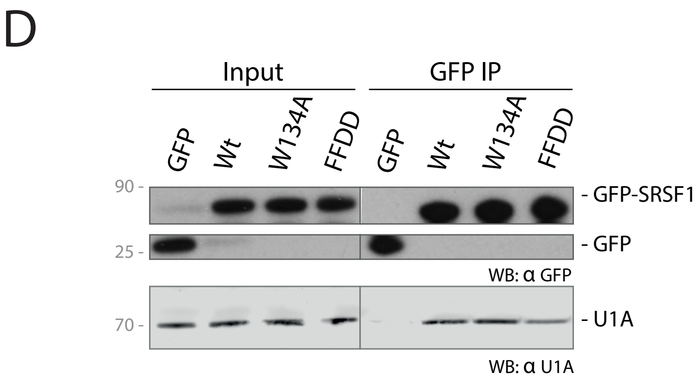

| Proteins Group Name | ID | GFP-SRSF1 wt / GFP (H/L) | ± SD | SRSF1 Known interactors according to BioGrid Previously detected by Akerman *et al*. (2015) | Nuclease resistant according Akerman *et al*. (2015) |
|---|---|---|---|---|---|
| Serine/arginine-rich splicing factor 1 | Q07955 | 15.27 | 2.43 | | |
| 40S ribosomal protein S12 | P25398 | 10.12 | 2.29 | yes | no |
| SRSF protein kinase 2;SRSF protein kinase 2 N-terminal;SRSF protein kinase 2 C-terminal | P78362;Q9UPE1 | 9.87 | 2.35 | | |
| Lupus La protein | P05455 | 9.85 | 0.80 | | |
| Histone H1x | Q92522 | 9.78 | 2.31 | yes | no |
| 40S ribosomal protein S24 | P62847 | 9.20 | 2.38 | | |
| 40S ribosomal protein S3a | P61247 | 8.39 | 2.49 | | |
| 40S ribosomal protein S9 | P46781 | 8.31 | 1.63 | | |
| 60S ribosomal protein L35 | P42766 | 8.07 | 1.74 | yes | no |
| 60S ribosomal protein L19 | P84098 | 7.83 | 1.52 | yes | no |
| 40S ribosomal protein S15 | P62841 | 7.52 | 1.67 | | |
| 40S ribosomal protein S4, X isoform | P62701;Q8TD47;P22090 | 7.41 | 1.24 | yes | no |
| Protein LTV1 homolog | Q96GA3 | 7.31 | 1.49 | | |
| Putative ATP-dependent RNA helicase DHX57 | Q6P158;Q7Z478 | 7.14 | 1.29 | yes | no |
| 60S ribosomal protein L17 | P18621 | 6.87 | 0.57 | yes | no |
| SRSF protein kinase 1 | Q96SB4 | 6.85 | 2.10 | | |
| 60S ribosomal protein L31 | P62899 | 6.79 | 1.56 | yes | yes |
| 40S ribosomal protein S6 | P62753 | 6.68 | 1.46 | yes | no |
| 60S ribosomal protein L10 | P27635;Q96L21 | 6.63 | 1.08 | | |
| Ribonucleases P/MRP protein subunit POP1 | Q99575 | 6.56 | 1.01 | | |
| 60S ribosomal protein L26-like 1;60S ribosomal protein L26 | Q9UNX3;P61254 | 6.53 | 1.30 | yes | no |
| 40S ribosomal protein S16 | P62249 | 6.49 | 1.38 | yes | no |
| 40S ribosomal protein S5;40S ribosomal protein S5, N-terminally processed | P46782 | 6.36 | 2.97 | yes | yes |
| 40S ribosomal protein S18 | P62269 | 6.26 | 1.05 | yes | no |
| 60S ribosomal protein L24 | P83731 | 6.24 | 1.59 | | |
| 40S ribosomal protein S11 | P62280 | 6.24 | 1.01 | | |
| 40S ribosomal protein S15a | P62244 | 6.21 | 1.51 | yes | no |
| 40S ribosomal protein S14 | P62263 | 6.12 | 1.15 | | |
| 40S ribosomal protein S8 | P62241 | 6.10 | 1.80 | | |
| Bystin | Q13895 | 6.09 | 1.19 | | |
| 40S ribosomal protein S2 | P15880 | 6.02 | 1.12 | | |
| U1 small nuclear ribonucleoprotein 70 kDa | P08621 | 5.98 | 2.22 | | |
| 40S ribosomal protein S13 | P62277 | 5.94 | 1.44 | | |
| 40S ribosomal protein S17 | P08708 | 5.78 | 1.47 | yes | no |
| 40S ribosomal protein S7 | P62081 | 5.70 | 0.96 | yes | yes |
| 60S ribosomal protein L23a | P62750 | 5.63 | 0.10 | yes | no |
| Heterogeneous nuclear ribonucleoprotein U-like protein 1 | Q9BUJ2 | 5.47 | 1.32 | yes | no |
| 60S acidic ribosomal protein P2 | P05387 | 5.41 | 0.61 | yes | no |

| | | | | | |
|---|---|---|---|---|---|
| THO complex subunit 4 | Q86V81 | 5.35 | 2.28 | yes | no |
| 40S ribosomal protein S28 | P62857 | 5.32 | 1.27 | yes | yes |
| Caprin-1 | Q14444 | 5.29 | 2.22 | yes | no |
| 40S ribosomal protein S26;Putative 40S ribosomal protein S26-like 1 | P62854;Q5JNZ5 | 5.22 | 0.87 | | |
| 60S ribosomal protein L28 | P46779 | 5.21 | 1.33 | yes | no |
| 40S ribosomal protein S23 | P62266 | 5.20 | 0.86 | | |
| 40S ribosomal protein S19 | P39019 | 5.17 | 1.66 | yes | no |
| 60S ribosomal protein L32 | P62910 | 5.11 | 1.05 | | |
| 60S ribosomal protein L8 | P62917 | 5.03 | 0.68 | yes | no |
| 28S ribosomal protein S17, mitochondrial | Q9Y2R5 | 4.95 | 0.02 | | |
| 60S ribosomal protein L21 | P46778 | 4.91 | 1.06 | | |
| 60S ribosomal protein L13a;Putative 60S ribosomal protein L13a protein RPL13AP3 | P40429;Q6NVV1 | 4.86 | 0.64 | yes | no |
| Nuclease-sensitive element-binding protein 1 | P67809;Q9Y2T7 | 4.77 | 1.60 | yes | no |
| Ras GTPase-activating protein-binding protein 2 | Q9UN86 | 4.73 | 1.96 | | |
| 60S acidic ribosomal protein P0;60S acidic ribosomal protein P0-like | P05388;Q8NHW5 | 4.63 | 0.58 | yes | no |
| Nucleolin | P19338 | 4.56 | 0.91 | yes | no |
| KRR1 small subunit processome component homolog | Q13601 | 4.54 | 2.36 | | |
| ATP-dependent RNA helicase DHX36 | Q9H2U1 | 4.54 | 1.15 | | |
| Cell division cycle and apoptosis regulator protein 1 | Q8IX12 | 4.50 | 1.04 | yes | yes |
| 60S ribosomal protein L15 | P61313 | 4.50 | 0.81 | | |
| Ras GTPase-activating protein-binding protein 1 | Q13283 | 4.49 | 2.06 | | |
| Zinc finger CCCH-type antiviral protein 1 | Q7Z2W4 | 4.47 | 0.78 | | |
| 60S ribosomal protein L12 | P30050 | 4.34 | 0.74 | yes | no |
| 40S ribosomal protein S10;Putative 40S ribosomal protein S10-like | P46783;Q9NQ39 | 4.34 | 1.19 | yes | no |
| La-related protein 1 | Q6PKG0 | 4.31 | 0.99 | yes | no |
| Heterogeneous nuclear ribonucleoprotein A/B | Q99729 | 4.29 | 2.02 | | |
| 28S ribosomal protein S23, mitochondrial | Q9Y3D9 | 4.28 | 0.52 | | |
| 60S ribosomal protein L13 | P26373 | 4.19 | 0.94 | | |
| Protein lin-28 homolog B | Q6ZN17 | 4.15 | 1.47 | | |
| 60S ribosomal protein L10a | P62906 | 4.15 | 0.53 | yes | no |
| 60S ribosomal protein L30 | P62888 | 4.15 | 0.91 | | |
| Insulin-like growth factor 2 mRNA-binding protein 3 | O00425 | 4.14 | 1.19 | | |
| 60S ribosomal protein L3 | P39023;Q92901 | 4.10 | 0.66 | | |
| 60S ribosomal protein L14 | P50914 | 4.06 | 0.68 | yes | no |
| 60S ribosomal protein L6 | Q02878 | 4.04 | 0.62 | yes | no |
| 60S ribosomal protein L27 | P61353 | 4.04 | 1.13 | yes | yes |
| 40S ribosomal protein S3 | P23396 | 4.03 | 0.70 | yes | no |
| 60S ribosomal protein L18a | Q02543 | 4.03 | 1.07 | yes | no |
| Heterogeneous nuclear ribonucleoprotein Q | O60506 | 4.02 | 1.73 | | |
| 60S ribosomal protein L7a | P62424 | 3.98 | 1.27 | yes | no |
| LINE-1 retrotransposable element ORF1 protein | Q9UN81 | 3.97 | 0.73 | | |
| Zinc finger CCHC domain-containing protein 3 | Q9NUD5 | 3.95 | 0.75 | | |
| 60S ribosomal protein L7 | P18124 | 3.95 | 0.62 | | |
| 60S ribosomal protein L36 | Q9Y3U8 | 3.95 | 1.10 | yes | no |

| | | | | | |
|---|---|---|---|---|---|
| 40S ribosomal protein S25 | P62851 | 3.92 | 0.16 | yes | no |
| Heterogeneous nuclear ribonucleoprotein A0 | Q13151 | 3.90 | 1.71 | | |
| 28S ribosomal protein S29, mitochondrial | P51398 | 3.86 | 0.77 | | |
| 28S ribosomal protein S27, mitochondrial | Q92552 | 3.86 | 1.58 | | |
| Periodic tryptophan protein 1 homolog | Q13610 | 3.84 | 0.89 | yes | yes |
| 40S ribosomal protein S20 | P60866 | 3.84 | 0.75 | | |
| 60S ribosomal protein L4 | P36578 | 3.82 | 0.58 | yes | no |
| 60S ribosomal protein L34 | P49207 | 3.82 | 0.59 | yes | no |
| 28S ribosomal protein S34, mitochondrial | P82930 | 3.81 | 0.48 | | |
| 28S ribosomal protein S22, mitochondrial | P82650 | 3.78 | 1.79 | yes | yes |
| Small nuclear ribonucleoprotein Sm D1 | P62314 | 3.76 | 1.55 | | |
| Ubiquitin carboxyl-terminal hydrolase 10 | Q14694 | 3.75 | 0.92 | | |
| Heterogeneous nuclear ribonucleoprotein U | Q00839 | 3.73 | 1.23 | | |
| Plasminogen activator inhibitor 1 RNA-binding protein | Q8NC51 | 3.73 | 0.35 | | |
| Small nuclear ribonucleoprotein Sm D2 | P62316 | 3.72 | 1.55 | | |
| Insulin-like growth factor 2 mRNA-binding protein 2 | Q9Y6M1 | 3.68 | 1.44 | yes | no |
| Probable ATP-dependent RNA helicase YTHDC2 | Q9H6S0 | 3.63 | 0.92 | yes | no |
| 60S ribosomal protein L35a | P18077 | 3.62 | 0.50 | yes | no |
| Putative ATP-dependent RNA helicase DHX30 | Q7L2E3 | 3.60 | 0.93 | yes | no |
| Transcriptional activator protein Pur-alpha | Q00577 | 3.59 | 1.32 | | |
| 60S ribosomal protein L18 | Q07020 | 3.58 | 0.80 | yes | no |
| Nucleolar RNA helicase 2 | Q9NR30 | 3.56 | 1.55 | | |
| Polyadenylate-binding protein 4 | Q13310;P0CB38 | 3.54 | 1.06 | yes | no |
| Double-stranded RNA-binding protein Staufen homolog 1 | O95793 | 3.54 | 0.55 | | |
| ATP-dependent RNA helicase A | Q08211 | 3.51 | 1.25 | yes | yes |
| Guanine nucleotide-binding protein-like 3 | Q9BVP2 | 3.50 | 1.02 | | |
| Small nuclear ribonucleoprotein Sm D3 | P62318 | 3.47 | 1.32 | | |
| Putative helicase MOV-10 | Q9HCE1 | 3.45 | 0.79 | | |
| N-acylneuraminate cytidylyltransferase | Q8NFW8 | 3.45 | 1.18 | | |
| Nucleolar GTP-binding protein 2 | Q13823 | 3.41 | 0.47 | | |
| Uncharacterized protein C7orf50 | Q9BRJ6 | 3.34 | 0.90 | | |
| 60S ribosomal protein L5 | P46777 | 3.33 | 1.44 | | |
| 60S ribosomal protein L9 | P32969 | 3.32 | 0.86 | yes | no |
| Insulin-like growth factor 2 mRNA-binding protein 1 | Q9NZI8 | 3.31 | 1.30 | | |
| ELAV-like protein 2;ELAV-like protein 4 | Q12926;P26378;Q14576 | 3.30 | 1.13 | | |
| Y-box-binding protein 3 | P16989 | 3.27 | 1.08 | yes | yes |
| Serine/arginine-rich splicing factor 7 | Q16629 | 3.23 | 1.10 | | |
| U4/U6 small nuclear ribonucleoprotein Prp3 | O43395 | 3.19 | 1.40 | | |
| Polyadenylate-binding protein 1 | P11940;Q9H361;Q4VXU2;Q96DU9 | 3.19 | 0.96 | yes | no |
| Interleukin enhancer-binding factor 2 | Q12905 | 3.18 | 1.04 | | |
| Constitutive coactivator of PPAR-gamma-like protein 1 | Q9NZB2;Q9NX05;Q5T035 | 3.18 | 0.68 | | |
| Heterogeneous nuclear ribonucleoprotein D0 | Q14103 | 3.16 | 1.19 | | |
| ELAV-like protein 1 | Q15717 | 3.09 | 1.09 | | |
| Pre-mRNA-processing factor 19 | Q9UMS4 | 3.08 | 0.97 | yes | no |

| | | | | | |
|---|---|---|---|---|---|
| Heterogeneous nuclear ribonucleoprotein A1;Heterogeneous nuclear ribonucleoprotein A1, N-terminally processed;Heterogeneous nuclear ribonucleoprotein A1-like 2 | P09651;Q32P51 | 3.07 | 1.31 | | |
| Interleukin enhancer-binding factor 3 | Q12906 | 3.05 | 1.19 | | |
| Nuclear fragile X mental retardation-interacting protein 2 | Q7Z417 | 2.99 | 0.71 | | |
| Protein regulator of cytokinesis 1 | O43663 | 2.97 | 0.65 | | |
| Small nuclear ribonucleoprotein-associated protein N;Small nuclear ribonucleoprotein-associated proteins B and B | P63162;P14678 | 2.90 | 0.55 | | |
| H/ACA ribonucleoprotein complex subunit 1 | Q9NY12 | 2.90 | 1.01 | | |
| Cell division cycle 5-like protein | Q99459 | 2.88 | 0.89 | yes | no |
| Serine/arginine-rich splicing factor 10 | O75494;Q8WXF0 | 2.88 | 0.43 | yes | yes |
| Nuclear cap-binding protein subunit 1 | Q09161 | 2.87 | 0.55 | | |
| 60S ribosomal protein L11 | P62913 | 2.87 | 0.28 | yes | no |
| 28S ribosomal protein S9, mitochondrial | P82933 | 2.86 | 1.05 | | |
| Nucleolar protein 16 | Q9Y3C1 | 2.85 | 0.82 | | |
| Regulator of nonsense transcripts 1 | Q92900 | 2.83 | 0.28 | | |
| Fragile X mental retardation syndrome-related protein 2 | P51116 | 2.81 | 0.61 | yes | yes |
| La-related protein 4 | Q71RC2 | 2.79 | 0.40 | | |
| Protein LSM12 homolog | Q3MHD2 | 2.77 | 0.62 | | |
| Fragile X mental retardation protein 1 | Q06787 | 2.73 | 0.77 | | |
| Interferon-inducible double-stranded RNA-dependent protein kinase activator A | O75569 | 2.69 | 0.18 | | |
| RNA-binding motif, single-stranded-interacting protein 1 | P29558;Q6XE24 | 2.67 | 0.67 | | |
| Replication factor C subunit 1 | P35251 | 2.66 | 0.56 | | |
| Fragile X mental retardation syndrome-related protein 1 | P51114 | 2.63 | 0.51 | yes | yes |
| Regulator of nonsense transcripts 3B | Q9BZI7 | 2.63 | 0.49 | | |
| Myb-binding protein 1A | Q9BQG0 | 2.58 | 0.57 | yes | no |
| RRP12-like protein | Q5JTH9 | 2.57 | 0.33 | | |
| Ataxin-2-like protein | Q8WWM7 | 2.51 | 0.63 | | |
| Probable ATP-dependent RNA helicase DDX5 | P17844 | 2.44 | 0.37 | yes | yes |
| RNA-binding protein Musashi homolog 2 | Q96DH6 | 2.43 | 0.43 | | |
| Kinesin-like protein KIF2A | O00139 | 2.42 | 0.04 | | |
| Luc7-like protein 3 | O95232 | 2.41 | 0.23 | yes | yes |
| 60S ribosomal protein L23 | P62829 | 2.37 | 0.23 | | |
| Transcription intermediary factor 1-beta | Q13263 | 2.37 | 0.29 | | |
| 60S ribosomal protein L22 | P35268 | 2.33 | 0.27 | | |
| Cold shock domain-containing protein E1 | O75534 | 2.30 | 0.30 | | |
| Translation machinery-associated protein 16 | Q96EY4 | 2.26 | 0.06 | | |
| Replication factor C subunit 4 | P35249 | 2.26 | 0.36 | | |
| Replication factor C subunit 3 | P40938 | 2.22 | 0.44 | | |
| Double-stranded RNA-binding protein Staufen homolog 2 | Q9NUL3 | 2.22 | 0.21 | | |
| AP-2 complex subunit beta;AP-1 complex subunit beta-1 | P63010;Q10567 | 2.18 | 0.06 | | |
| tRNA (cytosine(34)-C(5))-methyltransferase | Q08J23 | 2.15 | 0.16 | | |
| Pre-mRNA-splicing factor ATP-dependent RNA helicase DHX15 | O43143 | 2.15 | 0.43 | yes | yes |
| Glutamate-rich WD repeat-containing protein 1 | Q9BQ67 | 2.14 | 0.34 | | |
| Replication factor C subunit 2 | P35250 | 2.13 | 0.36 | | |
| Multiple myeloma tumor-associated protein 2 | Q9BU76 | 2.09 | 0.07 | | |

| | | | | | |
|---|---|---|---|---|---|
| Transcription factor A, mitochondrial | Q00059 | 2.09 | 0.33 | | |
| Probable ATP-dependent RNA helicase DDX17 | Q92841 | 2.05 | 0.29 | | |
| Serine/threonine-protein phosphatase PGAM5, mitochondrial | Q96HS1 | 2.04 | 0.11 | yes | yes |
| Poly(A) RNA polymerase, mitochondrial | Q9NVV4 | 2.03 | 0.09 | | |
| RNA-binding protein 6 | P78332 | 2.01 | 0.02 | | |
| Eukaryotic translation initiation factor 4 gamma 1 | Q04637 | 1.95 | 0.27 | | |
| Ataxin-2 | Q99700 | 1.94 | 0.31 | | |
| Heterogeneous nuclear ribonucleoprotein F;Heterogeneous nuclear ribonucleoprotein F, N-terminally processed | P52597 | 1.84 | 0.21 | yes | no |
| Leucine-rich PPR motif-containing protein, mitochondrial | P42704 | 1.81 | 0.16 | | |
| 40S ribosomal protein SA | P08865 | 1.81 | 0.19 | | |
| Protein PRRC2C | Q9Y520 | 1.75 | 0.06 | | |
| Cleavage and polyadenylation specificity factor subunit 1 | Q10570 | 1.75 | 0.15 | | |
| 40S ribosomal protein S21 | P63220 | 1.72 | 0.08 | | |
| Developmentally-regulated GTP-binding protein 1 | Q9Y295 | 1.71 | 0.05 | | |
| Cleavage and polyadenylation specificity factor subunit 3 | Q9UKF6 | 1.67 | 0.01 | | |
| Pumilio homolog 2 | Q8TB72 | 1.64 | 0.10 | | |
